# A photoswitchable HaloTag for spatiotemporal control of fluorescence in living cells

**DOI:** 10.1101/2024.12.18.629107

**Authors:** Franziska Walterspiel, Begoña Ugarte-Uribe, Jonas Weidenhausen, Anna Dimitriadi, Arif Ul Maula Khan, Christoph W. Müller, Claire Deo

## Abstract

Photosensitive fluorophores, which emission can be controlled using light, are essential for advanced biological imaging, enabling precise spatiotemporal tracking of molecular features, and facilitating super-resolution microscopy techniques. While irreversibly photoactivatable fluorophores are well established, reversible reporters which can be re-activated multiple times remain scarce, and only few have been applied in living cells using generalizable protein labelling methods. To address these limitations, we introduce chemigenetic photoswitchable fluorophores, leveraging the self-labelling HaloTag protein with fluorogenic rhodamine dye ligands. By incorporating a light-responsive protein domain into HaloTag, we engineer a tunable, photoswitchable HaloTag (psHaloTag), which can reversibly modulate the fluorescence of a bound dye-ligand via a light-induced conformational change. Our best performing psHaloTag variants show high performance *in vitro* and in living cells, with large, reversible, far-red fluorescence turn-on upon 450 nm illumination across various biomolecular targets. Together, this work establishes the chemigenetic approach as a versatile platform for the design of photoswitchable reporters, tunable through both genetic and synthetic modifications, with promising applications for dynamic imaging.

## INTRODUCTION

The ability to control the emission properties of a fluorophore using light is a powerful asset for biological imaging.^1, 2^ Indeed, the unique ability of photosensitive reporters to switch between a non-fluorescent and a fluorescent state under precise light conditions has enabled numerous biological applications, such as the marking and tracking of features of interest with exceptional spatiotemporal control,^3-5^ or the design of multifunctional tools such as light-gated biosensors.^6-8^ Importantly, photosensitive fluorophores are foundational to super-resolution microscopy, ^9^ for modalities including SMLM (single molecule localization microscopy),^1,10^ RESOLFT (reversible saturable optical linear fluorescence transition),^11-13^ and more recently MINFLUX (minimal photon fluxes),^14, 15^ enabling the visualization of biomolecules below the diffraction limit of light. Generally, photosensitive fluorophores can be categorized in two main groups, depending on whether they undergo an irreversible transformation upon illumination (i.e. photoactivatable, photoconvertible), or whether the transformation is reversible, either thermally or upon illumination with light at a different wavelength (i.e. photoswitchable, photochromic). While many irreversible systems, based either on fluorescent proteins (FPs) or synthetic dye scaffolds, have been developed, photoswitchable systems which can be re-activated multiple times are lacking. To date, most photoswitchable reporters applied in biological samples are based on fluorescent proteins.^12, 16-22^ However, these inherit the limitations of conventional FPs, with generally low brightness and photostability, particularly in the far-red region of the spectrum, high pH sensitivity and oxygen requirement. Accordingly, synthetic photoswitchable fluorophores are highly attractive, but their engineering has been notoriously difficult. Indeed, Förster resonance energy transfer-based photoswitchable fluorophores generally display low contrast between the OFF and ON forms, and can present stability issues due to differential bleaching of the donor and acceptor.^23-25^ Intrinsically fluorescent photoswitches such as diarylethenes and spiropyrans show coupled switching and excitation, which limits brightness or switching efficiency.^26-28^ A promising approach is the modulation of the electronic conjugation of workhorse fluorophores, as shown in rhodamine-lactams,^29-32^ in which the open-close equilibrium can be indirectly affected by light. However, most of these systems require low wavelength light (≤ 405 nm) for activation, and their use in living cells with broadly applicable protein labelling methods remains scarce. Together, this highlights the need for novel approaches and mechanisms for fluorescence photocontrol, combining high brightness, efficient photoswitching using visible light and live cell compatibility.

To address these challenges, we introduce a “chemigenetic” approach for photoswitchable fluorescent systems, leveraging the HaloTag self-labeling protein and bright fluorogenic rhodamine ligands. Fluorogenic rhodamines predominantly exist in a closed, non-fluorescent state in solution. ^33^ Upon binding to the HaloTag protein,^34, 35^ the change in dye environment shifts the equilibrium toward the open, fluorescent form, resulting in a large fluorescence turn-on. This property has been recently exploited for the design of biosensors, based on genetically encoded sensing motifs fused to HaloTag in which an analyte-dependent conformational change alters the equilibrium and hence the fluorescence of the dye.^36-39^ Here, we repurpose this approach to engineer a photoswitchable fluorescent system by integrating a light-responsive protein domain into the HaloTag protein (Figure 1). Upon illumination, the conformational change of the protein photoswitch alters the dye environment, shifting its equilibrium toward the open, fluorescent state. This led to the development of a tunable, photoswitchable HaloTag (psHaloTag), which can modulate the fluorescence of its bound ligand in response to light. This system retains the excellent photophysical properties of established rhodamines, with entirely electronically decoupled photoswitching and fluorescence processes. Advantageously, the fluorogenic dye becomes switchable only when bound to the protein tag, guaranteeing low background from unbound reporters. Our best performing psHaloTag variants exhibit robust far-red fluorescence turn-on *in vitro* as well as in living cells across various subcellular targets, demonstrating the potential of the chemigenetic approach for fluorescence photocontrol in complex biological samples.

**Figure 1.**
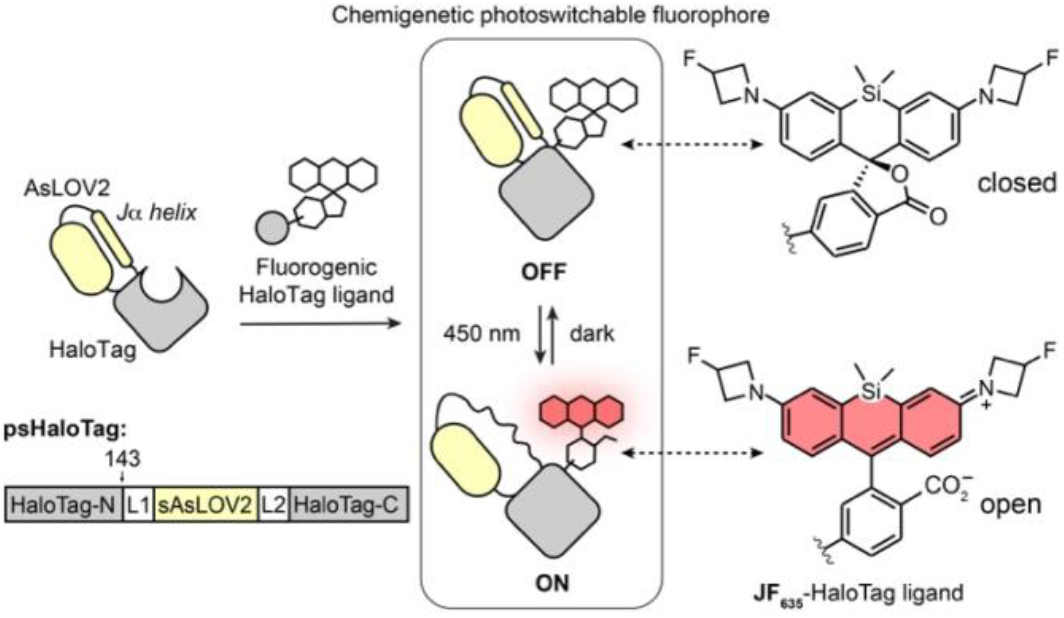
General principle of photoswitchable HaloTag (psHaloTag) and open-closed equilibrium of **JF**_**635**_**-HaloTag ligand** (HTL). L1 and L2 denote linkers.

## RESULTS AND DISCUSSION

### Engineering of a photoswitchable HaloTag

To engineer a photoswitchable self-labelling tag, we set out to introduce a photoswitchable domain into the HaloTag, in close proximity to the dye binding site. We reasoned that the incorporation of the light-sensing motif into HaloTag would change its interaction with the fluorogenic dye ligand, resulting in low fluorescence in the dark. Upon illumination, the conformational change of the photo-responsive domain would subsequently alter the environment of the bound fluorogenic dye, resulting in a shift in open-close equilibrium and concomitant fluorescence increase. As the genetically-encoded photoswitch, we selected the light-oxygen-voltage 2 domain of *Avena sativa* phototropin 1 (AsLOV2).^40, 41^ This small 16.5 kDa protein uses flavin mononucleotide (FMN) as a cofactor, and comprises two terminal α-helices, the small N-terminal A’α and the large C-terminal Jα helix. Exposure to 450 nm light induces the formation of a metastable photo-adduct between a cysteine side chain and the FMN cofactor, which leads to the undocking and unfolding of the Jα helix. This large conformational change has been extensively used to control the activity of effector molecules, and is fundamental for the design of optogenetic tools.^40^ The photo-reaction is reversible, with backfolding to the thermally stable state occurring spontaneously in the dark.^42^

For our design, we used a truncated version of the AsLOV2 domain between residues 408 and 543, termed sAsLOV2, previously shown to be functional with improved conformational coupling when fused to nanobodies,^43,44^ and added a glycine to each terminus of the domain to provide a slight flexibility. Using the crystal structure of HaloTag bound to tetramethylrhodamine (PDB: 6Y7A),^45^ six residues on the surface of the HaloTag protein (143, 154, 160,166, 178, and 180, FigureS1a) were selected for engineering due to their proximity to the position of the bound fluorophore ligand. Accordingly, a series of 16 HaloTag-sAsLOV2 chimeras were generated, either by insertion of sAsLOV2 into HaloTag, or fusion of sAsLOV2 to the termini of a circularly permuted HaloTag (FigureS1b). We selected **JF**_**635**_**-HTL** (λ_max_/λ_em_ = 635/652 nm) as the fluorophore ligand for engineering, due to its large turn-on in far-red absorbance and fluorescence upon binding to HaloTag (∼113 fold),^46^ and the success of this dye ligand for engineering sensitive chemigenetic biosensors based on conformational change.^36-39^ The different constructs were evaluated in bacterial lysates (Figure S2), and we assessed their ability to bind **JF**_**635**_**-HTL**, and their far-red fluorescence intensity, in the dark (Figure S1c,d). Among them, only constructs with HaloTag modified at positions 143 and 154 retained efficient binding to **JF**_**635**_**-HTL** (i.e. near complete binding after 2 hours of incubation at room temperature). This is in agreement with previous work, in which these HaloTag sequence regions have also been shown to be successfully targeted for domain insertion or circular permutation.^36, 47, 48^ Insertion of a SG linker between the HaloTag and LOV domains to add flexibility led to faster binding kinetics for some of the slower-binding constructs, however still not reaching complete binding after 2 hours (e.g. at position 180, FigureS1d). The constructs generated from modification at position 154 showed a basal fluorescence in the dark comparable to that of HaloTag labelled with **JF**_**635**_**-HTL**, while constructs generated from modifications at position 143 showed 8−35% of HaloTag fluorescence, indicating that the fluorophore is only partially open in those constructs. We then assessed the light-response of the efficiently-binding constructs, by measuring the change in far-red absorption upon photoswitching of the LOV domain, elicited by illumination at 450 nm using a custom-made LED device (Figure S3). Among them, only 3 constructs showed a change in absorption, all with a small decrease upon illumination at 450 nm, which was fully reversible when incubated in the dark (FigureS1e). Indeed, the insertion of sAsLOV2 after residues 143 (**#2**, Figure S1) or 154 (**#3**) of HaloTag led to A/A_0_ of 0.5 and 0.7 respectively. The terminal fusion of sAsLOV2 to HaloTag circularly permuted at the same positions did not lead to any absorption change. This could be explained by the fact that only one terminus of the LOV domain is attached to the tag in those constructs, resulting in limited conformational coupling. Surprisingly however, construct **#1** with the sAsLOV2 fused directly at the N-terminus of the unmodified HaloTag protein showed a small light response, with a 10% decrease in **JF**_**635**_ absorption upon illumination. This might be due to the relative orientation of the two domains in this construct, which could bring them in close proximity to interact, as previously observed with EGFP-HaloTag fusions.^49^ Based on this initial screening, we selected ins143HaloTag-sAsLOV2 (construct **#2**) for further engineering, showing the largest change in absorption, and advantageously a lower absorption in the dark state which suggests substantial room for engineering towards large turn-ons.

To facilitate screening of a large number of mutants, we set out to slow down the thermal relaxation of the photo-reaction. Indeed, the original, “wild-type” AsLOV2 displays a t_1/2_ ∼ 1 min, too fast for robust measurements of the ON state properties using steady-state UV-Vis and fluorescence spectroscopies, and we therefore introduced a point mutation known to slow-down the thermal relaxation.^42, 50^ The V416L (numbered according to the AsLOV2 sequence) mutation led to an unstable protein, however V416I led to a construct displaying efficient photoswitching, with a t_1/2_ ∼ 6.8 min at room temperature. This “slow-relaxing” construct, ins143HaloTag-sAsLOV2(V416I), was called psHaloTag0.1, and used as the starting point for improvement.

Several rounds of protein engineering were then performed (Figure 2a, Figure S4), with screening in *E. coli* lysates to quantify ligand binding efficiency, fluorescence turn-on in response to light, and associated photoswitching kinetics (Figure S2). To reverse the response of the system, and therefore lead to a preferable fluorescence turn-on upon illumination at 450 nm, we first investigated the effect of linker length between HaloTag and sAsLOV2, with a two amino acid linker extension on either side based on the HaloTag sequence (either FA at the N-term, or PE at the C-term of the sAsLOV2 domain, respectively, Figure S4). These insertions had no measurable effect on the photoswitching and fluorescence properties, and we therefore performed site-saturation mutagenesis of selected residues within those linkers. Screening in bacterial lysate led to the identification of psHaloTag0.1-G283P (termed psHaloTag0.2), which showed a small **JF**_**635**_ fluorescence turn-on upon illumination. To further increase the turn-on, we next targeted residues in proximity to the fluorophore binding site, expected to be critical for function (20 amino acids, Figure S4), and performed site-saturation mutagenesis. We identified mutation E143W, leading to a large improvement of fluorescence turn-on upon illumination (psHaloTag0.3). An additional round of targeted site-saturation mutagenesis in residues around the HaloTag surface (11 amino acids) led to the identification of two mutants A285W (psHaloTag1a, F/F_0_ = 4.6) and A285L (psHaloTag1b, F/F_0_ = 3.5, Figure 2b). Gratifyingly, both constructs showed full reversibility in the dark, and several ON/OFF cycles could be performed (Figure 2c). While psHaloTag1b showed smaller turn-on, the photoswitching kinetics were substantially faster than psHaloTag1a, warranting its selection for further investigation (Figure 2d).

**Figure 2.**
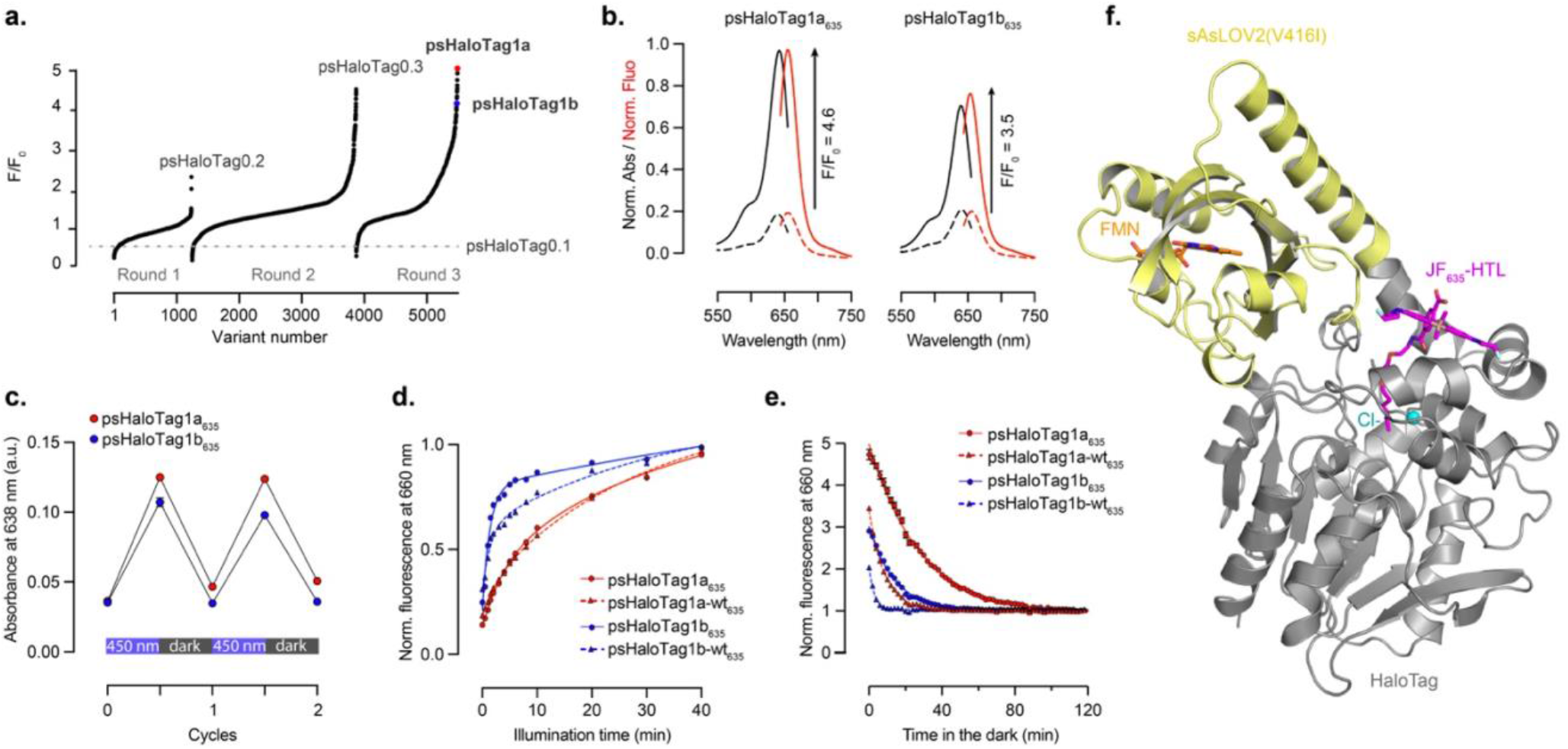
Engineering and characterization of psHaloTag. **a**. Sequential rounds of site-saturation mutagenesis with selected mutants highlighted in each round. **b**. Normalized absorption (black) and fluorescence (red) spectra of psHaloTag1a_635_ and psHaloTag1b_635_ in the dark (dashed lines) and following illumination at 450 nm (solid lines). **c**. Absorbance at 638 nm of psHaloTag1a_635_ and psHaloTag1b_635_ during two cycles of illumination at 450 nm followed by incubation in the dark at room temperature. **d**. Kinetics of the fluorescence turn-on of **JF**_**635**_ upon illumination at 450 nm for the psHaloTag constructs. **e**. Kinetics of the fluorescence turn-off of **JF**_**635**_ upon incubation at room temperature in the dark. **f**. Crystal structure of psHaloTag1a labelled with **JF**_**635**_**-HTL** in the dark state (PDB:9HKF).

### Characterization of psHaloTag in vitro

psHaloTag1a, psHaloTag1b and the hits isolated in each round of mutagenesis (Figure S4, Table S2) were fully characterized as purified proteins after labelling with **JF**_**635**_**-HTL** (the protein-dye conjugates will be hereafter referred to as psHaloTag_635_). We measured their absorption and fluorescence spectra in the dark and after illumination at 450 nm, and recorded the kinetics of the photoswitching (turn-on) and thermal relaxation (turn-off) at room temperature in the dark (Table 1, Figure S5). While psHaloTag0.1_635_ showed an absorption-driven change in fluorescence (A/A_0_ = 0.4, F/F_0_ = 0.5), psHaloTag0.2_635_ displayed a small absorption turn-off and fluorescence turn-on (A/A_0_ = 0.95, F/F_0_ = 1.1). This suggests that, in this construct, the fluorescence modulation stems predominantly from a change in fluorescence quantum yield, albeit small, and not a shift in the open-close equilibrium of the dye. Introduction of mutation E143W in psHaloTag0.3_635_ resulted in very different properties, recovering an equilibrium-driven mechanism (A/A_0_ = 3.1, F/F_0_ = 3.4). This was retained in improved mutants psHaloTag1a_635_ (A/A_0_ = 4.7, F/F_0_ = 4.6) and psHaloTag1b_635_ (A/A_0_ = 3.1, F/F_0_ = 3.5), isolated from the third round of mutagenesis. We confirmed that psHaloTag1a retained fast binding to **JF**_**635**_**-HTL** in the dark, with labelling complete in 15 minutes in the experimental conditions (Figure S6), and that psHaloTag1a_635_ showed low pH sensitivity over the range of pH 6−9 (Figure S7). Both psHaloTag1a_635_ and psHaloTag1b_635_ showed high Φ_F_ > 0.68, and displayed ∼40% of the brightness of HaloTag_635_ in their ON states due to lower extinction coefficients, which demonstrates that the dye does not fully open in those constructs. While further improvements are certainly possible to shift the dye further towards the open form in the ON state, it is worth noting that psHaloTag1a_635_ is substantially brighter than other red-emitting photoswitchable proteins, e.g. 3-fold brighter than PSmOrange (λ_max_/λ_em_ = 634/662 nm)^19^ and rsFusionred1 (λ_max_/λ_em_ = 577/605 nm),^13^ and 16-fold brighter than rsCherry(λ_max_/λ_em_ = 572/610 nm).^21^

**Table 1.** Sequence information and properties of the psHaloTag variants labelled with **JF**_**635**_**-HTL**. Wavelengths, extinction coefficients and quantum yields correspond to the ON state of the constructs, measured after illumination at 450 nm. t_1/2, relax_ corresponds to the half-life of the thermal relaxation of the **JF**_**635**_ signal (measured in the dark at room temperature). n.m.: not measured.

| Construct | L1 | L2 | mutations | $\lambda_{\text{max}}/\lambda_{\text{em}}$ (nm) | $\epsilon$ (M<sup>-1</sup>.cm<sup>-1</sup>) | $\Phi_{\text{Fl}}$ | A/A<sub>0</sub> | F/F<sub>0</sub> | $t_{1/2, \text{relax}}$ (min) |
| --- | --- | --- | --- | --- | --- | --- | --- | --- | --- |
| HaloTag | / | / | / | 640/656 | 81000 | 0.75 | 0 | 0 | / |
| psHaloTag0.1 | G | G | / | 641/656 | 12800 | 0.70 | 0.4 | 0.5 | 6.8 |
| psHaloTag0.2 | FAG | P | / | 641/656 | 46300 | 0.75 | 0.95 | 1.1 | n.m. |
| psHaloTag0.3 | FAG | P | E143W | 640/654 | 30900 | 0.76 | 3.1 | 3.4 | 12.6 |
| psHaloTag1a | FAG | P | E143W, A285W | 642/655 | 38200 | 0.68 | 4.7 | 4.6 | 19.5 |
| psHaloTag1a-wt | FAG | P | E143W, A285W, I155V | 641/656 | 25200 | n.m. | 3.6 | 4.4 | 5.9 |
| psHaloTag1b | FAG | P | E143W, A285L | 639/655 | 30200 | 0.74 | 3.1 | 3.5 | 9.7 |
| psHaloTag1b-wt | FAG | P | E143W, A285L, I155V | 639/655 | 20900 | n.m. | 2.4 | 3.1 | 1.8 |

In order to assess the composition of the ON state of psHaloTag1a_635_ and psHaloTag1b_635_, we generated “dark” and “lit” mimics of these constructs, introducing mutations shown to lock AsLOV2 in either the folded, or unfolded states (“dark mimic”: C189A, “lit mimic”: I271E, A275E in psHaloTag1a and psHaloTag1b, Figure S8).^51, 52^ Spectroscopic evaluation revealed that the dark mimic closely resembles the dark state of the parent constructs. The lit mimic shows slightly higher intensity in both absorption and fluorescence, with a small hypsochromic shift. While using these “mimics” to replicate the two extreme cases of the photocycle constitutes an approximation, these results suggest that the OFF state corresponds to ∼100% of folded proteins, and the ON state contains >85% of unfolded proteins for both psHaloTag1a and psHaloTag1b, therefore supporting efficient photoswitching.

We then examined the photoswitching kinetics of the different constructs, measuring the absorption and fluorescence of the FMN cofactor (λ_max_/λ_em_ = 450/525 nm) and **JF**_**635**_ (λ_max_/λ_em_ = 640/660 nm) (Figure S5, Table S2). Under illumination at 450 nm, the FMN absorption decreases, due to the formation of the covalent bond with the cysteine residue in the LOV scaffold, while the absorption of **JF**_**635**_ increases as the Jα helix unfolds, which elicits a conformational change in the HaloTag protein close to the dye. The LOV-FMN adduct formation occurred rapidly in all constructs, reaching a photostationary state after the first illumination. In contrast, the **JF**_**635**_ turn-on was substantially slower. This could be explained as the far-red fluorescence modulation arises from a conformational change which needs to be propagated from the Jα helix to the HaloTag, and likely requires important rearrangements, while outcompeting the thermal relaxation process. psHaloTag1b_635_ showed ∼6-fold faster turn-on of the far-red signal than psHaloTag1a_635_. Similar observations were made for the thermal relaxation kinetics where psHaloTag1a_635_ and 1b_635_ showed similar t_1/2,relax_(FMN) of 6.7 and 7.9 min, but distinct far-red kinetics with t_1/2,relax_(JF_635_) of 19.5 and 9.7 min, respectively. The difference could be explained by the presence of the additional bulky tryptophane W285 at the hinge region in construct 1a, which might slow-down conformational rearrangement. Interestingly, the initial negative variant psHaloTag0.1_635_ showed a different behavior, with similar t_1/2,relax_ for both the FMN and **JF**_**635**_ signals. These differences suggest that the conformational rearrangement of the protein scaffold is the rate limiting step in the fluorescence turn-on of the system. To investigate this further, we reversed the initial “slow” mutation introduced in psHaloTag0.1 for screening purposes, leading to the “wild-type” psHaloTag1a-wt (psHaloTag1a(I155V), with I155V in the psHaloTag sequence numbering corresponding to I416V in AsLOV2 sequence numbering) and psHaloTag1b-wt (psHaloTag1b(I155V)) mutants. Both retained fluorescence turn-on, albeit slightly smaller than the parent constructs with F/F_0_ of 4.4 and 3.1 for psHaloTag1a-wt and psHaloTag1b-wt, respectively. This could be explained by the competing faster thermal relaxation (Table 1, Table S2). Indeed, the t_1/2,ON_(JF_635_) were unchanged compared to the parent constructs, but the turn-off kinetics were, as expected, substantially faster, with t_1/2_,_relax_(FMN) = 0.6 min for both constructs and t_1/2_,_relax_(JF_635_) = 5.9 and 1.8 min for psHaloTag1a-wt_635_ and psHaloTag1b-wt_635_, respectively (Figure 2d,e). While the far-red fluorescence modulation remains slower than the LOV domain photocycle itself, this demonstrates that faster kinetics can be achieved, with minimal impact on the ON/OFF contrast.

### Tunability by varying the dye ligand

A key property of chemigenetic systems is the high tunability afforded by modifying the dye ligand. ^53^ Indeed, using a different fluorogenic HaloTag ligand can lead to a broad range of spectral properties, dynamic ranges and kinetics, while generally retaining function. We therefore tested a panel of dyes in combination with psHaloTag1a, including fluorogenic Janelia Fluor (JF) dyes and Max-Planck (MaP) dyes (Figure S9).^36, 46, 54-56^ All dyes tested led to a fluorescence turn-on upon illumination, albeit to different extent. The MaP dyes as well as the O-, C- and P-bridged rhodamines led to smaller turn-ons, with F/F_0_ ≤ 2.0, whereas the other Si-rhodamines, closely resembling **JF**_**635**_, led to substantially higher turn-ons. In this dye series (compounds **1-5**, Figure 3, Figure S10), the trend followed the electron withdrawing capability of the azetidine substituents, and hence the open-close equilibrium of the dyes, with dyes more closed generally resulting in lower fluorescence intensities in both the ON and OFF states. Both **JF**_**629**_**-HTL** and **JF**_**630**_**-HTL** led to larger fluorescence turn-ons of 6.7 and 9.4 folds, respectively. Among compounds **1-5**, the kinetics were strongly dye-dependent, with larger ON/OFF fluorescence ratio generally correlating with slower turn-on kinetics, while the relaxation kinetics were varied with no particular trend. These results demonstrate that the psHaloTag system can be finely tuned by changing the dye ligand, with adjustable brightness, turn-on ratio and kinetics, while retaining the advantageous far-red spectral features.

**Figure 3.**
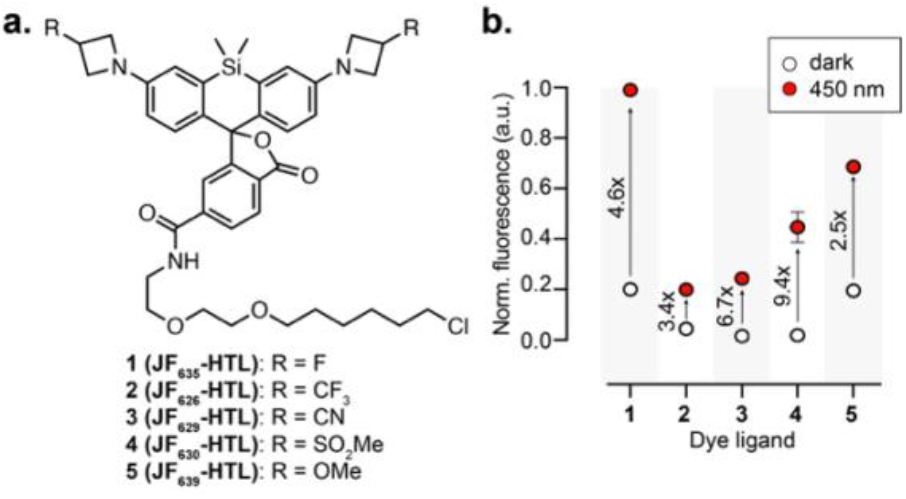
**a**. Chemical structure of **JF**_**635**_**-HTL** derivatives. **b**. Normalized fluorescence in the OFF (white dots) and ON (red dots) states for psHaloTag1a labelled with dyes **1-5**. Numbers on the graph indicate F/F_0_. Fluorescence was normalized to the ON state of psHaloTag1a_635_.

### Structure of psHaloTag

In order to gain insights into the mechanism of psHaloTag, we used AlphaFold for structure prediction.^57, 58^ The predicted structures of psHaloTag1a and psHaloTag1b indicate that the Jα helix is fused to one of the helices of HaloTag, resulting in an elongated helix (Figure S11). This could be responsible for the mechanism of the system, where, upon illumination, unfolding of the Jα helix would lead to partial unfolding of the HaloTag helix itself, propagating the conformational changes to residues in close proximity to the fluorophore ligand, with concomitant shift of the equilibrium towards the open form. The predicted structures of psHaloTag1a and 1b appear similar, with a slightly different bent in the extended helix. However, AlphaFold predicts uncertainty in these linker regions, which makes it difficult to unambiguously conclude on structural differences between the two constructs. In addition, structure prediction does not provide information on the position and state of the dye ligand or FMN cofactor. We therefore additionally solved the crystal structure of psHaloTag1a bound to **JF**_**635**_**-HTL**, which diffracted to 2.4 Å resolution (PDB: 9HKF), and which allowed us to place the ligands in the electron density. The crystals formed in space group C121 and contained two biological assemblies (psHaloTag1a, FMN, Cl^-^, and **JF**_**635**_**-HTL**) per asymmetric unit. The structure was determined by molecular replacement using a HaloTag crystal structure (PDB: 5Y2Y) as an initial model followed by several cycles of building and refining. The final structure also contains eleven molecules glycerol which was used as cryo-protectant during data collection. The two psHaloTag1a molecules adapt a near-identical conformation with a root-mean-square deviation (RMSD) of 0.33 Å (for 431 Cα residues) (Figure S12a,b) The extended J*α*helix is in a folded state in both molecules and consists of about 1/3 HaloTag and 2/3 LOV domain residues. Our experimentally determined structure is also in agreement with the AlphaFold prediction with a RMSD of 1.64 Å (for 371 Cα residues) (Figure S12c). The major difference between the structure and the prediction is attributed to a slightly different bend of the J*α* helix, resulting in a shifted orientation of the LOV domain. The FMN cofactor, **JF**_**635**_**-HTL** fluorophore ligands and coordinated Cl^-^ ions were clearly visible in the electron density. Based on the density, both FMN molecules appear to be unbound to the adjacent C189 residues, and the folded helix of the LOV domain suggests crystallization of the OFF, dark state of the protein (Figure S13). **JF**_**635**_**-HTL** adopted the same position as similar fluorophore ligands on HaloTag (Figure S13b-d), and while the electron density around the dyes does not allow for unambiguous identification of the open or closed conformation, the open conformation resulted in the best fit (Figure S12d). While we would expect a primarily closed conformation in solution for the OFF state, the observed open form could be explained by the tight crystal packing around the fluorophore (Figure S12a) as the surface-exposed dyes from the two chains face each other in the center of the asymmetric unit. The two performance-increasing mutations E143W and A285W are in proximity to each other, and A285W to the headgroup of the **JF**_**635**_**-HTL** fluorophore. These tryptophane residues could form an extended hydrophobic network together with neighboring hydrophobic residues (I150, L147, and additional residues towards the center of the protein) which could improve the stabilization/interaction of the fluorophore and improve the performance. Overall, both the predicted and the crystal structures, show that the HaloTag and LOV domains are tightly arranged and connected by a long helix, which ensures efficient propagation of the conformational change to the HaloTag in close proximity to the fluorophore.

### Characterization in living cells

Finally, we set out to investigate the performance of psHaloTag1a in living cells, to confirm that photoswitching was retained in the native cellular environment, and that the system was broadly usable across various subcellular locations. U2OS cells expressing psHaloTag1a-T2A-EGFP targeted to the nucleus (H2B), mitochondria (TOMM20) or F-actin (LifeAct) were stained with **JF**_**635**_**-HTL** (1 µM, 2 h), washed, and imaged using widefield microscopy (Figure 4). Specific labelling of psHaloTag1a was confirmed by the dim far-red fluorescence in the OFF state and, gratifyingly, a large fluorescence turn-on was observed for all targets after illumination at 450 nm for 3 minutes, while the HaloTag control showed no fluorescence change after illumination (Figure 4a-g). The dynamic range in cells was very consistent for the three subcellular targets evaluated, with F/F_0_ between 3.5−4.1 for psHaloTag1a_635_ (Figure 4g), and about 30% of the brightness of HaloTag_635_ in cells, in excellent alignment with the *in vitro* measurements (Figure 4h). As expected, **JF**_**630**_**-HTL** led to a significantly larger turn-on, with F/F_0_ = 7.0 for LifeAct-psHaloTag1a_630_ (Figure S14). In the dark, the system was fully reversible in living cells with both dyes tested, and illumination cycles could be successfully repeated (Figure 4i).

**Figure 4.**
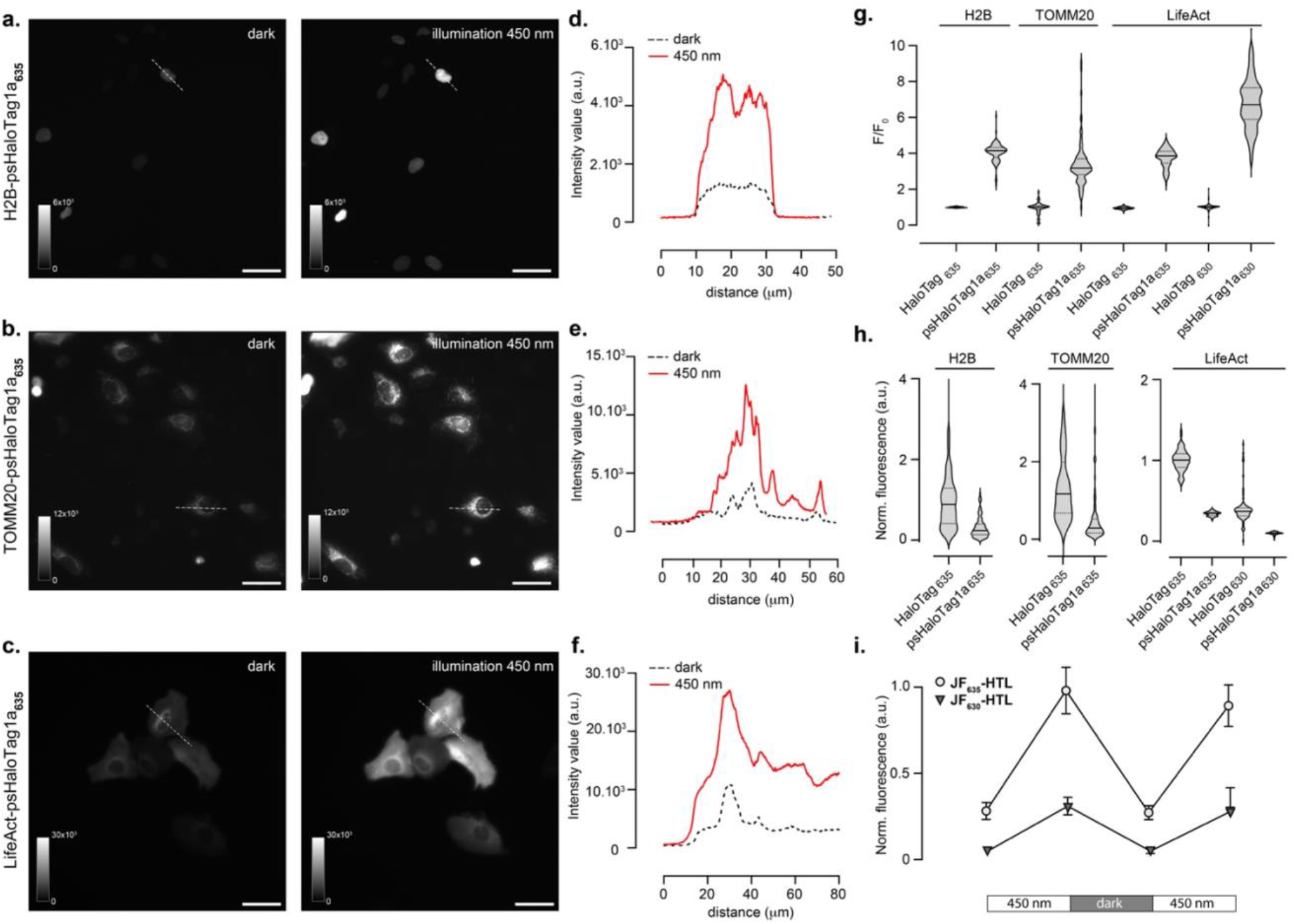
Characterization of psHaloTag1a in living cells. **a-c**. Representative images of U2OS cells expressing psHaloTag1a-T2A-EGFP targeted to the nucleus (**a**.), mitochondria (**b**.) or actin (**c**.) and labelled with **JF**_**635**_**-HTL**, in the dark and after illumination at 450 nm. Scale bars: 50 μm. **d-f**. Intensity line profiles corresponding to the images in **a-c**., respectively. **g**. F/F_0_ for HaloTag and psHaloTag1a targeted to different cellular locations. **h**. Normalized fluorescence intensity for the same cells as in **g**. Fluorescence intensity is the ratio of far-red to EGFP fluorescence, and was normalized to the value of HaloTag_635_ for each respective localization. **i**. Normalized fluorescence intensity of cells expressing LifeAct-psHaloTag1a-T2A-EGFP and labelled with either **JF**_**635**_**-HTL** or **JF**_**630**_**-HTL**, upon cycles of illumination at 450 nm followed incubation in the dark. Quantification in panels **g-i**. was performed on N > 100 cells for each condition.

## CONCLUSION

In this work, we developed a photoswitchable HaloTag (psHaloTag) that enables light-controlled reversible fluorescence modulation of a far-red rhodamine fluorophore through a light-driven conformational change. We systematically explored the insertion of a light-sensitive AsLOV2 domain into the HaloTag protein, and improved the performance of the system through rationally guided site-saturation mutagenesis. The psHaloTag platform leverages the superior properties of synthetic dyes such as their high brightness, while conferring them a light-responsive behavior. The system was largely tunable, and dynamic range, photoswitching and thermal relaxation kinetics could be finely modified by introducing point mutations or varying the dye ligand. The best performing psHaloTag-fluorophore combination shows close to 10-fold fluorescence increase *in vitro*, and is substantially brighter than existing red-shifted photoswitchable protein. We demonstrated that this system works robustly in living cells across multiple subcellular targets, showing reversible far-red fluorescence turn-on. Looking forward, we expect that continued directed evolution and further structure-based engineering of this platform will further enhance fluorescence turn-on ratios. Furthermore, the current iteration of psHaloTag is exclusively thermally reversible, which imposes constraints on imaging speed, and limitations for applications requiring rapid reversibility. Future iterations of the platform could incorporate optically reversible light-sensitive domains, enabling more versatile spatiotemporal control. Together, this work represents a step forward in integrating synthetic dyes and protein scaffolds for fluorescence photoswitching. By further improving the dynamic range and enhancing switching mechanisms, this approach has the potential to open new avenues in dynamic imaging and super-resolution microscopy.

## SUPPLEMENTARY MATERIAL

Supplementary figures and tables, materials and methods (pdf).

The crystal structure of psHaloTag1a labelled with **JF**_**635**_**-HTL** has been deposited to the Protein Data Bank (PDB: 9HKF).

The image analysis code used for segmentation and intensity quantification along with test data can be found at (CBA/psHaloTag-mitochondria-intensity-quantification·GitLab).

## ACKNOWLEDGMENTS

This work was supported by the European Molecular Biology Laboratory (EMBL) and the Chan Zuckerberg Initiative (Deep Tissue Imaging grants no. 2020-225346 and no. 2024-337799).

The authors acknowledge: The Johnsson group (MPI for Medical Research) for sharing fluorophores and plasmids; The Lavis group (Janelia Research Campus, HHMI) for sharing fluorophores, and Luke Lavis for contributive discussions; Federico Marotta (EMBL) for help implementing AlphaFold/ColabFold); Lucia Alvarez for help with kinetics measurements; Anmol Gautam and the ESRF synchrotron (Grenoble, France) staff for support in using beamline ID23-2; The Core Facilities at EMBL: Electrical and Mechanical Workshops (LED illumination device); Karine Lapouge and the Protein Expression and Purification Core Facility (protein purification, characterizations); Manuel Gunkel, Marko Lampe and the Advanced Light Microscopy Facility (live cell imaging).

## ABBREVIATIONS

FMN: flavin mononucleotide
FP: fluorescent protein
HTL: HaloTag ligand
LOV: light-oxygen-voltage
ps: photoswitchable

