## Supplementary Information for "A photoswitchable HaloTag for spatiotemporal control of fluorescence in living cells"

### Supplementary Figures

**Figure S1.** Engineering of psHaloTag0.1. **a**. Crystal structure of HaloTag bound to tetramethylrhodamine HaloTag ligand (PDB: 6Y7A).[1] The yellow spheres indicate positions selected for introduction of the sAsLOV2 domain. **b**. Schematic representation of the different types of constructs generated including insertion of sAsLOV2 into HaloTag, and N- or C-terminal fusion of sAsLOV2 to circularly permutated HaloTag. “XXX” in the name of the constructs denotes the amino acid position of the insertion or circular permutation. White boxes in the scheme depict the different linkers (G, glycine; cpL, circular permutation linker $(GGTGGS)_{x3}$) **c**. Performance of the HaloTag-sAsLOV2 constructs tested, showing the normalized fluorescence in the dark vs the bound fraction of **$JF_{635}$-HTL** after 2 hours of incubation. Values were normalized to $HaloTag_{635}$. While screening in lysate leads to high variability in measurements, constructs could be classified between “non-binding”, “slow binding” or “fast binding”, and among the fast-binding constructs, separated whether the **$JF_{635}$** fluorophore is mostly open or closed. The red-colored dots indicate the three constructs that showed a fluorescence change upon illumination at 450 nm (#1, #2, #3). **d.** Characterization of the generated HaloTag-sAsLOV2 proteins generated labelled with **$JF_{635}$-HTL**. The fluorescence intensity, binding capability, and far-red absorption change upon illumination at 450 nm were measured. **e**. Absorption spectra of constructs #1, #2, #3 labelled with **$JF_{635}$-HTL** (100 nM), measured as purified protein in the dark and after illumination at 450 nm. n.m: not measured.

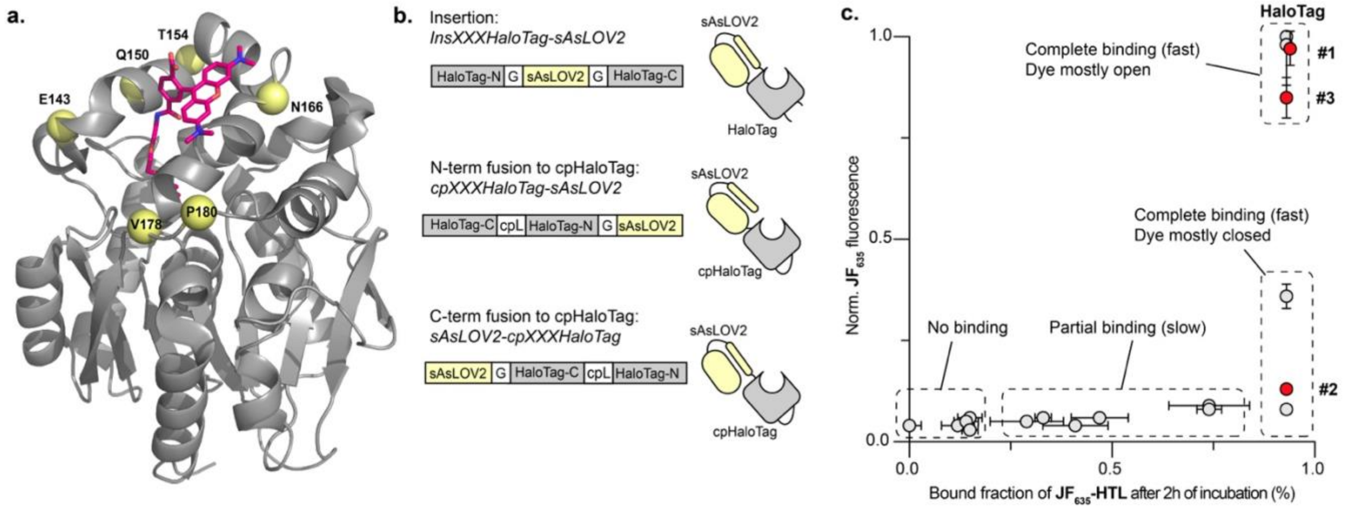

d.

| Position | # | Construct | Norm. $JF_{635}$ fluorescence | Bound fraction of $JF_{635}$-HTL (%) | $A/A_0$ |
|---|---|---|---|---|---|
| NA | | HaloTag | 1 | 93 ± 1 | n.m. |
| NA | | sAsLOV2 | 0.05 ± 0.01 | 14 ± 3 | n.m. |
| NA | #1 | sAsLOV2-HaloTag | 0.97 ± 0.04 | 94 ± 1 | 0.9 |
| 143 | #2 | ins143HaloTag-sAsLOV2 | 0.13 ± 0.01 | 93 ± 1 | 0.5 |
| 143 | | sAsLOV2-cp143HaloTag | 0.36 ± 0.03 | 93 ± 1 | 0 |
| 143 | | cp143HaloTag-sAsLOV2 | 0.08 ± 0.01 | 93 ± 1 | 0 |
| 150 | | ins150HaloTag-sAsLOV2 | 0.04 ± 0.01 | 12 ± 4 | n.m. |
| 154 | #3 | ins154HaloTag-sAsLOV2 | 0.85 ± 0.05 | 93 ± 1 | 0.7 |
| 154 | | sAsLOV2-cp154HaloTag | 0.98 ± 0.1 | 93 ± 1 | 0 |
| 166 | | ins166HaloTag-sAsLOV2 | 0.03 ± 0.01 | 15 ± 2 | n.m. |
| 166 | | sAsLOV2-cp166HaloTag | 0.04 ± 0.01 | 0 ± 3 | n.m. |
| 178 | | ins178HaloTag-sAsLOV2 | 0.06 ± 0.01 | 15 ± 3 | n.m. |
| 178 | | sAsLOV2-cp178HaloTag | 0.06 ± 0.01 | 33 ± 2 | n.m. |
| 178 | | cp178HaloTag-sAsLOV2 | 0.04 ± 0.01 | 41 ± 8 | 0 |
| 180 | | ins180HaloTag-sAsLOV2 | 0.06 ± 0.01 | 47 ± 7 | 0 |
| 180 | | sAsLOV2-cp180HaloTag | 0.05 ± 0.01 | 29 ± 9 | n.m. |
| 180 | | ins180Halo-SG-sAsLOV2 | 0.09 ± 0.01 | 74 ± 10 | 0 |
| 180 | | ins180Halo-sAsLOV2-SG | 0.08 ± 0.01 | 74 ± 3 | 0 |

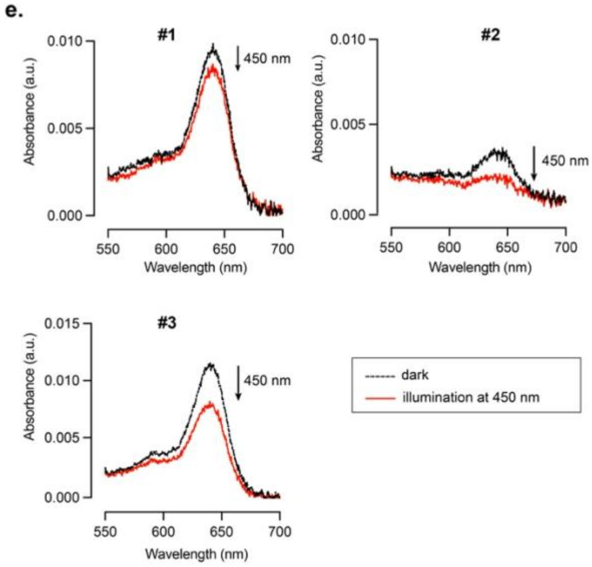

**Figure S2.** Schematic of the *in vitro* screening assay used for the engineering of psHaloTag. Following generation of a plasmid library through site-saturation mutagenesis and transformation in *E. coli*, individual colonies were manually picked in a 96 well-plate. After expression and lysis as described in the methods, **$JF_{635}$-HTL** (50 nM) was added to each well and incubated for 2 hours at room temperature in the dark. The labelled lysate was divided into two. Half was used to measure the binding efficiency, by addition of excess purified HaloTag protein to capture all unbound **$JF_{635}$-HTL**, and calculate the bound-fraction using fluorescence measurements. The second half of the lysate was used to perform the photoswitching assay, by three cycles of illumination at 450 nm (3 minutes for each cycle). Fluorescence measurements enabled to calculate $F/F_0$ of each construct, and additionally assess the photoswitching kinetics. r.t.: room temperature.

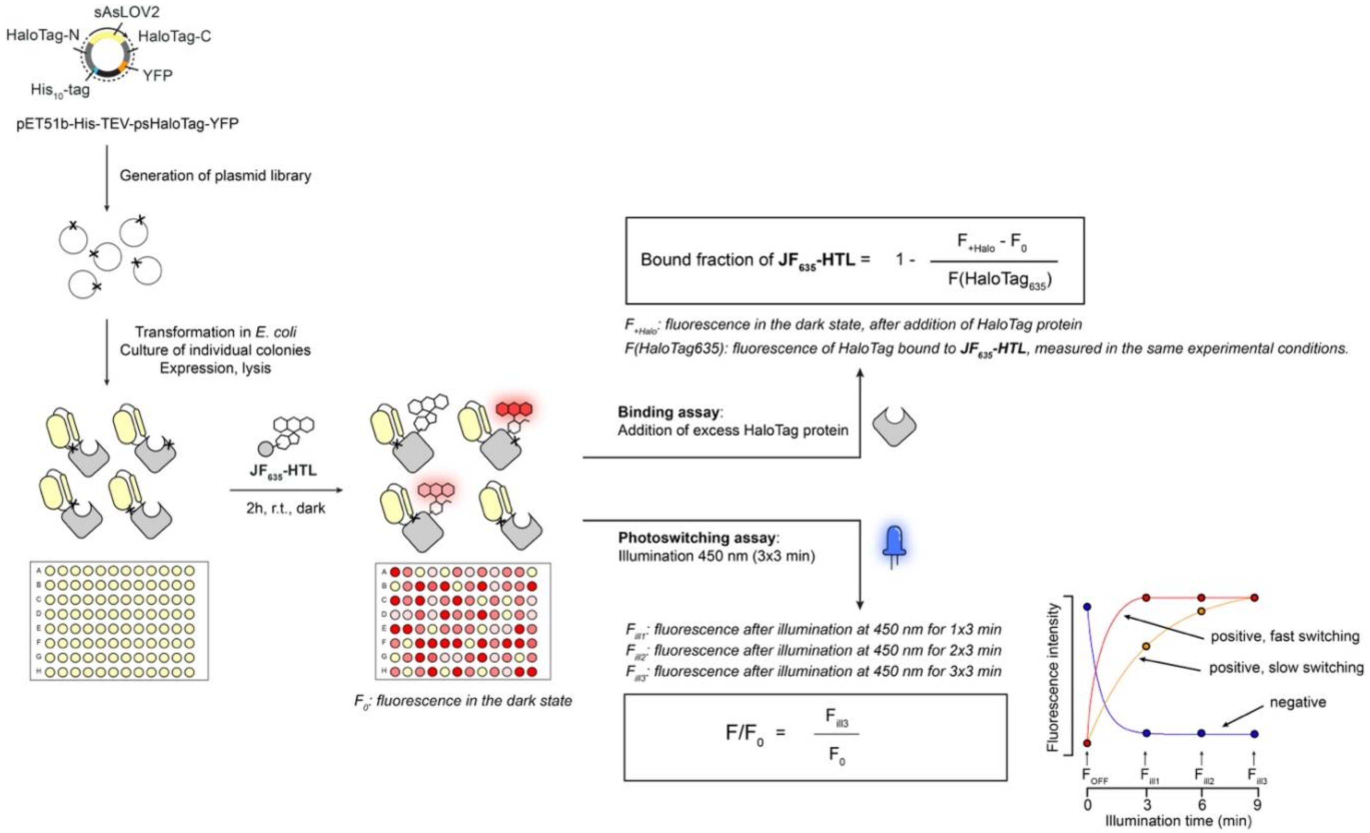

**Figure S3**. Picture of the custom-built illumination device used for inducing photoswitching of psHaloTag *in vitro* and *in cellulo*. The illumination box was made to fit a 96-well plate, and comprises 24 LEDs at 450 nm, and a controller to trigger illumination.

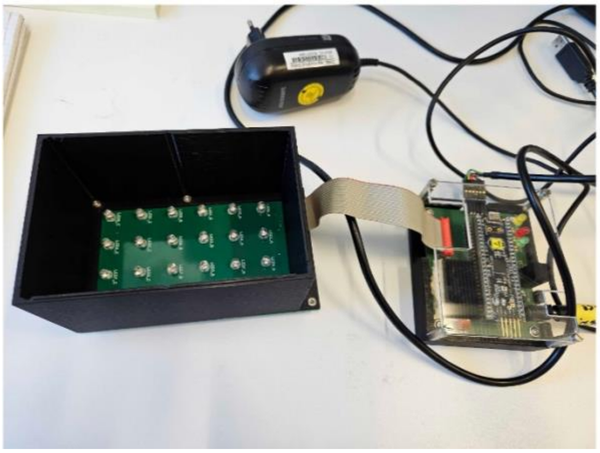

**Figure S4.** Stepwise engineering of psHaloTag1a and 1b, showing detail of the positions saturated in each round and schematic sequence representation of isolated variants. The first round of mutagenesis was performed in parallel on 3 constructs: psHaloTag0.1, psHaloTag0.1 with insertion of PE in the linker at the C-side of sAsLOV2, and psHaloTag0.1 with insertion of FA in the linker at the N-side of sAsLOV2. Full sequences of the isolated constructs in each round can be found in Table S1.

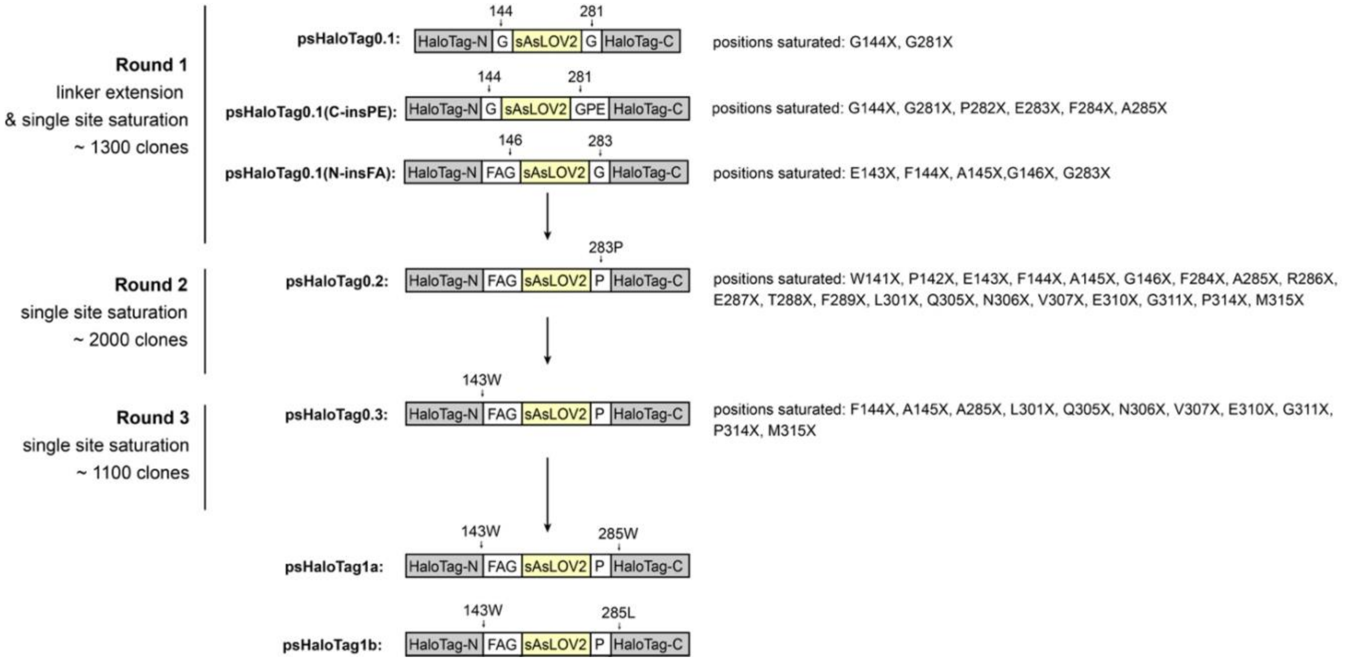

**Figure S5**. Absorption (left panel) and emission (center left panel) spectra of selected psHaloTag variants labelled with **$JF_{635}$-HTL**, in the dark (dashed black) and after illumination at 450 nm (red). Kinetics of the photoswitching (turn-on under 450 nm illumination, center right panel) and thermal relaxation (turn-off in the dark, right panel) of the FMN cofactor (yellow) and **$JF_{635}$** (red) of selected variants. Absorption measurements were performed at 5 μM dye ligand and 7.5 μM protein, and fluorescence measurements at 1 μM dye ligand and 1.5 μM protein, in triplicate. Kinetic profiles are plotted as mean and SEM, and fitted to a single exponential curve. n.m.: not measured.

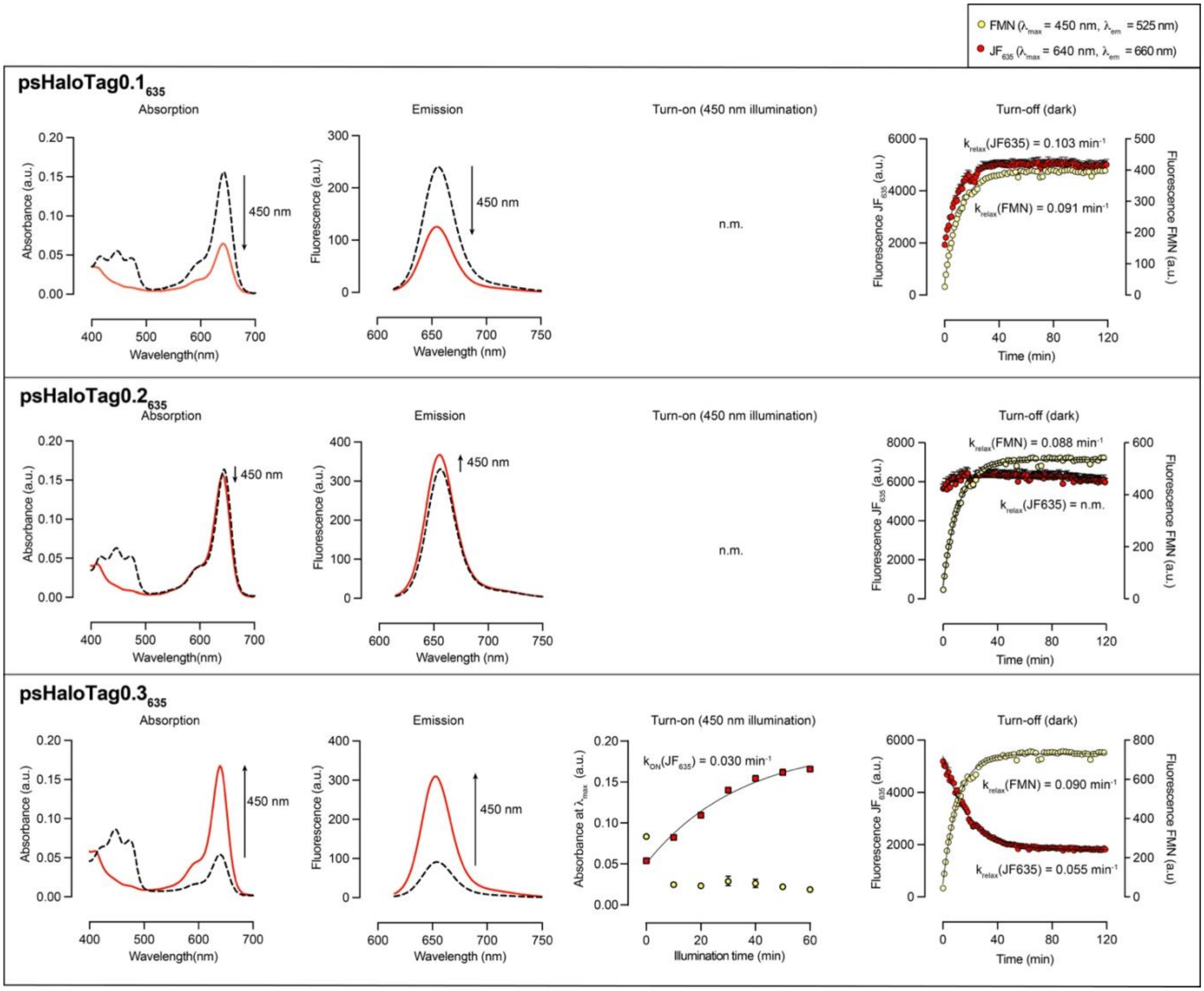

**Figure S5** – Continued.

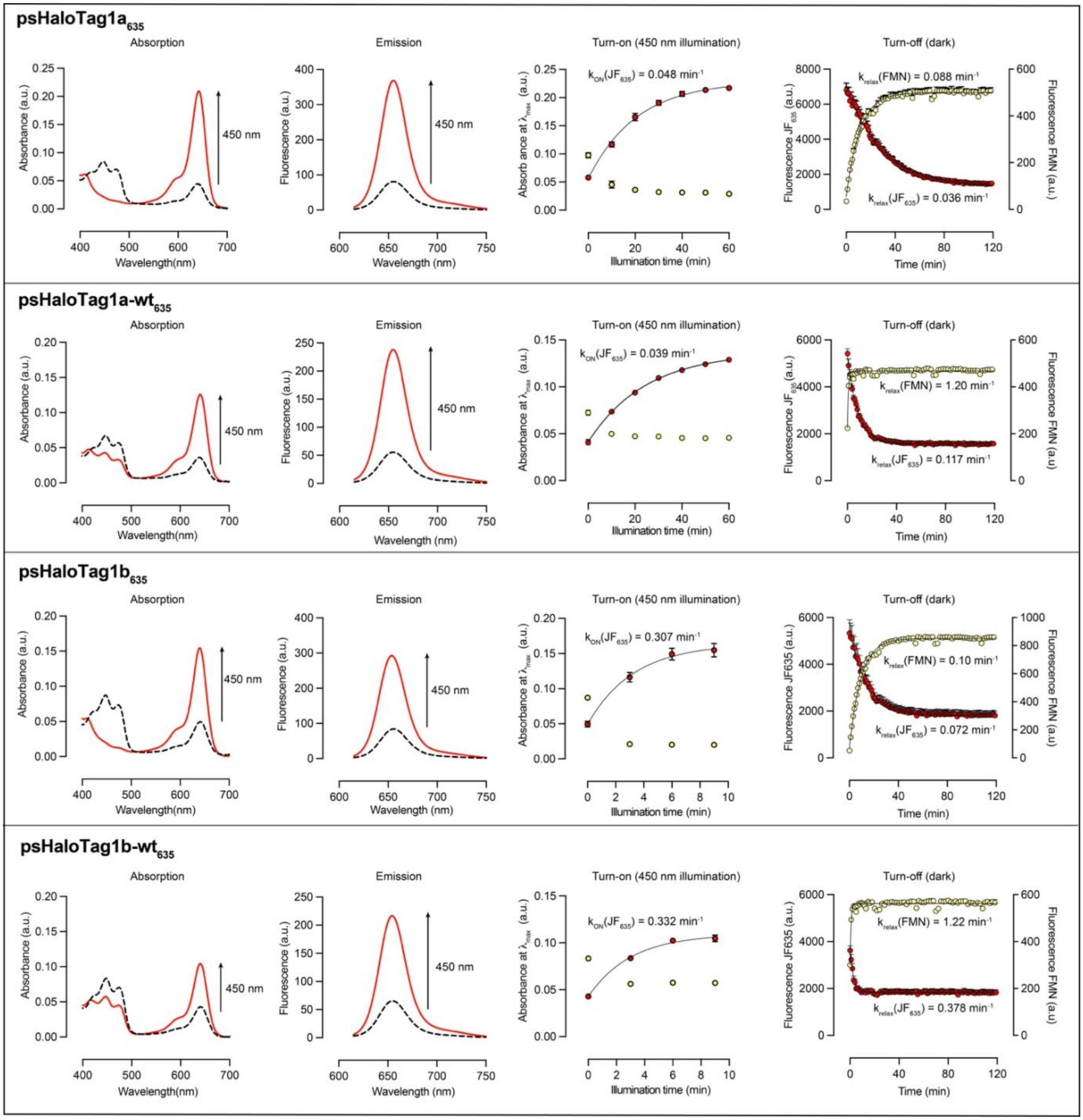

**Figure S6.** Labelling of psHaloTag1a (7.5 µM) with **JF$_{635}$-HTL** (5 µM) measured by the increase in absorbance at 640 nm when incubated in the dark. Labelling was complete after 15 minutes. Mean and SEM for 3 replicates.

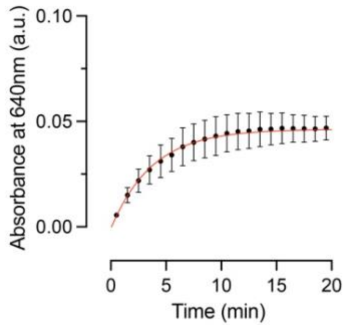

**Figure S7.** pH-dependence of the fluorescence and photoswitching properties of psHaloTag1a$_{635}$. **a**. Fluorescence intensity ($\lambda_{ex}/\lambda_{em}$ = 640/660 nm) of psHaloTag1a$_{635}$ at different pHs in the dark and after illumination at 450 nm. **b**. F/F$_0$ as a function of pH for psHaloTag1a$_{635}$. **c**. Apparent relaxation rate constant at room temperature in the dark for the FMN signal and the **JF$_{635}$** signal for psHaloTag1a$_{635}$. Plots represent mean and SEM for 3 replicates.

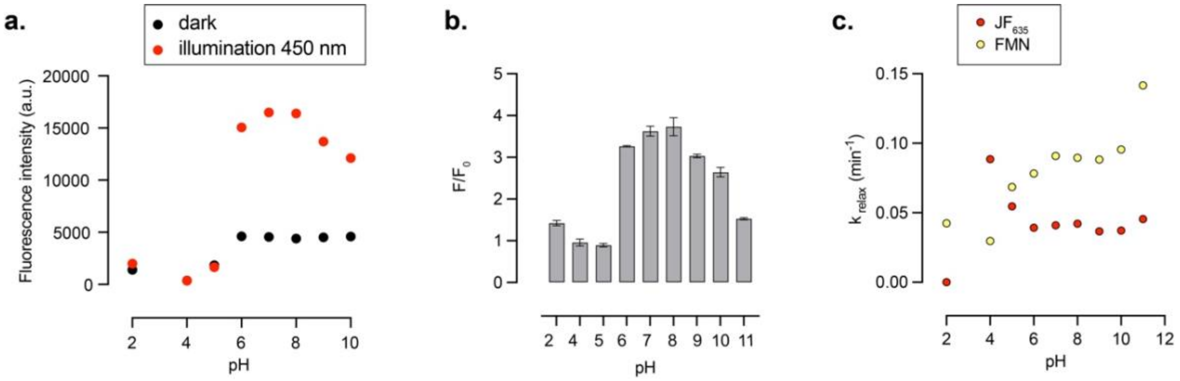

**Figure S8**. Absorption and fluorescence spectra of psHaloTag1a$_{635}$, psHaloTag1b$_{635}$ and psHaloTag1a$_{630}$ in the dark (solid black line) and following illumination at 450 nm (solid red line). Absorption and emission spectra of the “dark” mimic (mutant C189A, dashed black line) and “lit” mimic (double mutant I271E, A275E, dashed red line) for the same constructs labelled with the same fluorophore ligands.

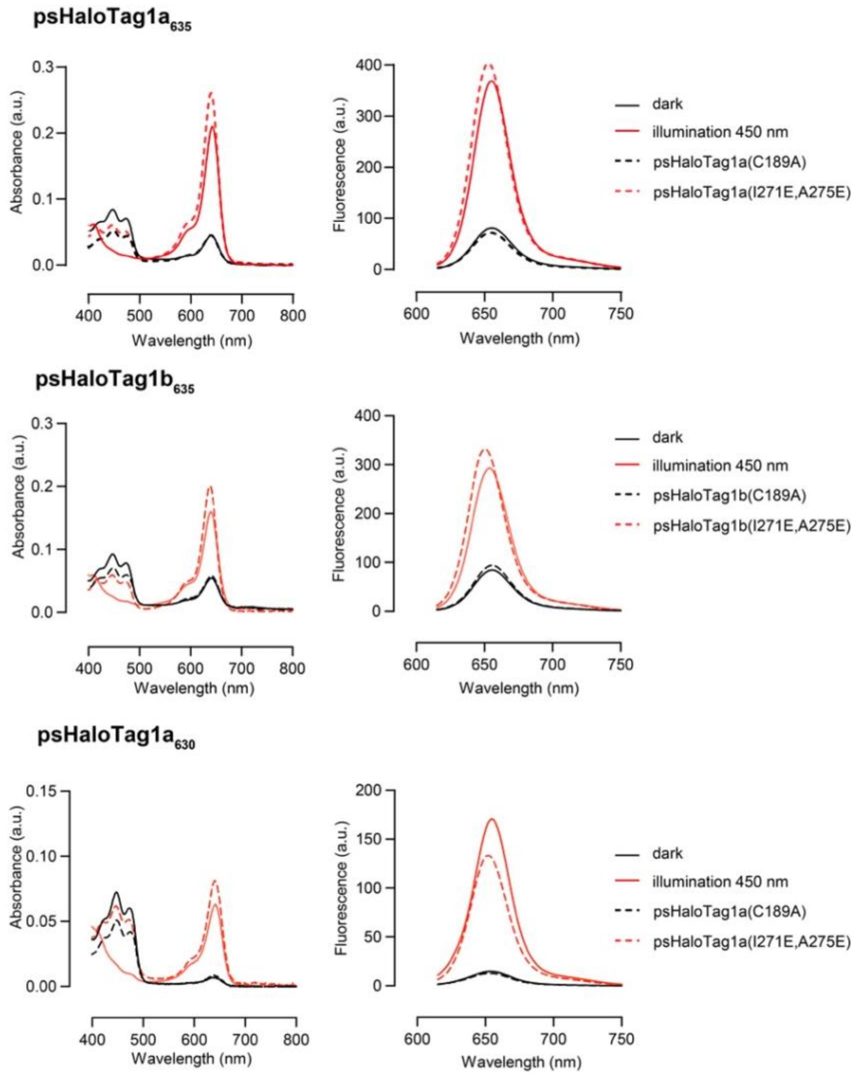

**Figure S9. a**. General structure and open close equilibrium of fluorogenic HaloTag ligands. **b**. Chemical structure of fluorogenic HaloTag ligands tested, and properties of psHaloTag1a when labelled with those dye ligands. n.m.: not measured.

**a.**

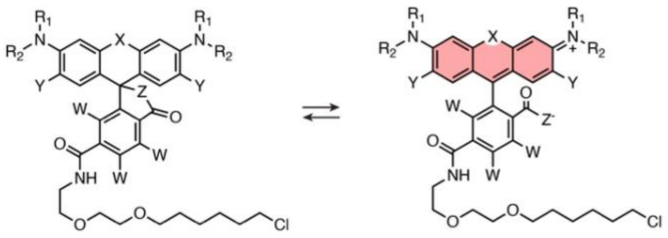

**b.**

| | X | $R_1$-N-$R_2$ | Y | Z | W | $\lambda_{max}/\lambda_{em}$ (nm) | $A/A_0$ | $F/F_0$ | $t_{1/2, ON}(JF_{635})$ (min) | $t_{1/2, relax}$(FMN) (min) | $t_{1/2, relax}(JF_{635})$ (min) |
|---|---|---|---|---|---|---|---|---|---|---|---|
| $JF_{525}$-HTL | O | N (azetidine) F, F | H | O | H | 525/549 | n.m | 1.2 | n.m. | n.m. | n.m. |
| $JF_{526}$-HTL | O | N (azetidine) F, F | F | O | H | 526/550 | n.m. | 1.2 | n.m. | n.m. | n.m. |
| $JF_{585}$-HTL | C | N (azetidine) F, F | H | O | H | 585/609 | n.m. | 2.0 | n.m. | n.m. | n.m. |
| $MaP_{618}$-HTL | C | N (dimethyl) | H | N-$SO_2NMe_2$ | H | 618/635 | n.m. | 1.2 | n.m. | n.m. | n.m. |
| $JF_{626}$-HTL (2) | Si | N (azetidine) $CF_3$ | H | O | H | 626/638 | 2.1 | 3.4 | 5.5 | 9.8 | 23.1 |
| $JF_{629}$-HTL (3) | Si | N (azetidine) CN | H | O | H | 629/648 | 4.0 | 6.7 | 11.3 | 10.5 | 16.1 |
| $JF_{630}$-HTL (4) | Si | N (azetidine) $SO_2Me$ | H | O | H | 630/649 | 9.2 | 9.4 | 28.9 | 13.3 | 26.6 |
| $JF_{635}$-HTL (1) | Si | N (azetidine) F | H | O | H | 635/652 | 4.7 | 4.6 | 14.3 | 7.9 | 19.5 |
| $JF_{639}$-HTL (5) | Si | N (azetidine) OMe | H | O | H | 639/656 | n.m. | 2.5 | n.m. | n.m. | n.m. |
| $MaP_{700}$-HTL | Si | N (fused ring) | | N-CN | H | 700/720 | n.m. | 1.6 | n.m. | n.m. | n.m. |
| $JF_{711}$-HTL | P(O)Ph | N (azetidine) F | H | O | F | 711/732 | n.m. | 1.2 | n.m. | n.m. | n.m. |

**$JF_{635}$-HTL** was synthesized as previously described.[2] The HaloTag ligands of **$JF_{525}$,**[2] **$JF_{526}$,**[3] **$JF_{585}$,**[2] **$JF_{626}$,**[4] **$JF_{629}$,**[4] **$JF_{630}$,**[4] **$JF_{639}$,**[4] **$JF_{711}$**[5] were provided by the Lavis group (Janelia Research Campus, HHMI). The HaloTag ligands of **$MaP_{618}$**[6] and **$MaP_{700}$**[6] were provided by the Johnsson group (Max Planck Institute for Medical Research).

**Figure S10**. Absorption (left panel) and emission (center left panel) spectra psHaloTag1a labelled with **$JF_{626}$-HTL**, **$JF_{629}$-HTL** or **$JF_{630}$-HTL** in the dark (dashed black) and after illumination at 450 nm (red). Absorption kinetics measured at $\lambda_{max}$ of the photoswitching (turn-on under 450 nm illumination, center right panel) and thermal relaxation (turn-off in the dark, right panel) for the signal of the FMN cofactor (yellow) and dye (red) of selected variants. Absorption measurements were performed at 5 μM dye ligand and 7.5 μM protein, and fluorescence measurements at 1 μM dye ligand and 1.5 μM protein, in triplicate. Kinetics curves are plotted as mean and SEM, and fitted to a single exponential curve. $\lambda_{max}$ = 637 nm for **$JF_{626}$**; 638 nm for **$JF_{629}$;** 640 nm for **$JF_{630}$**; 450 nm for FMN.

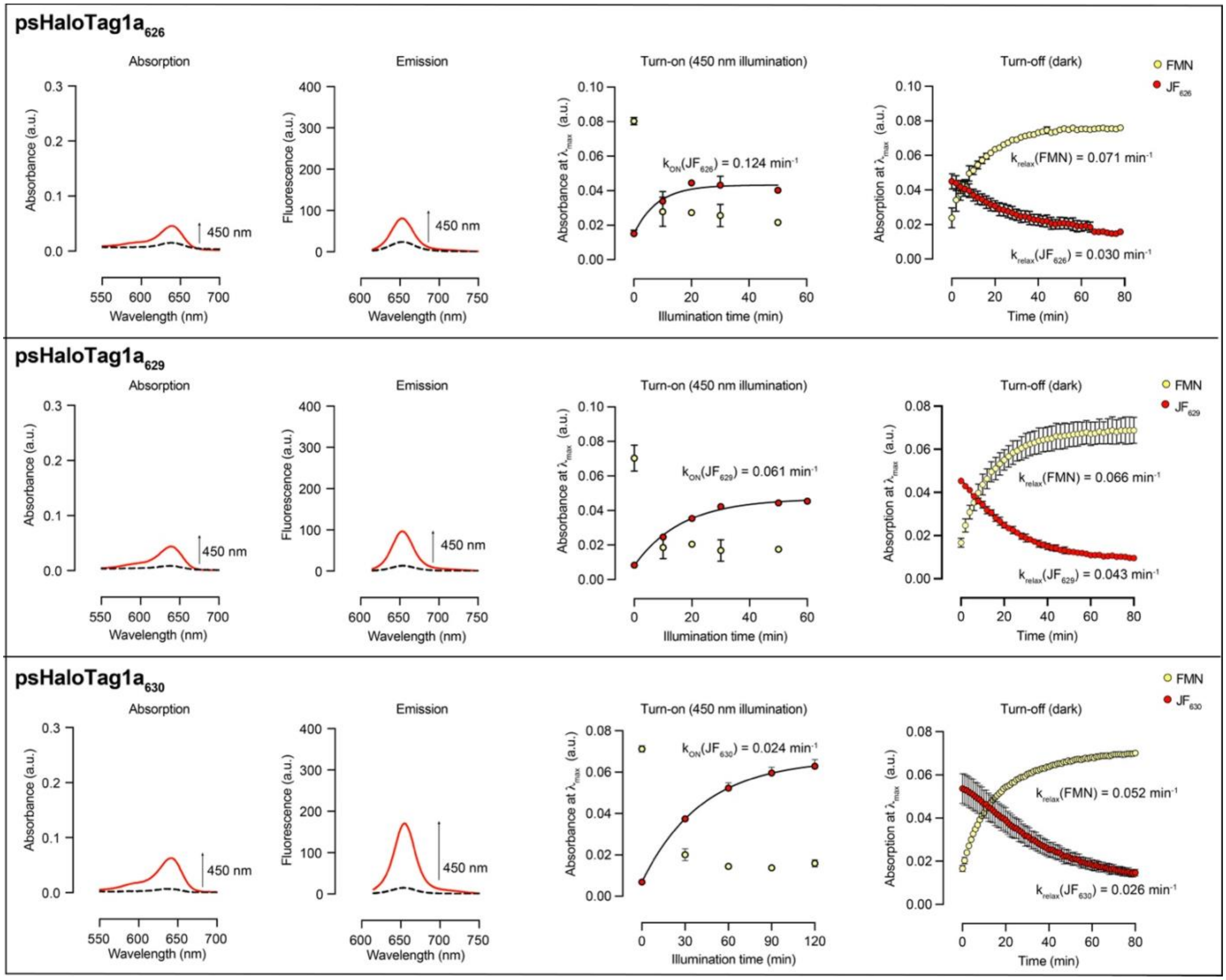

**Figure S11**. AlphaFold predicted structures (**a,c**) and corresponding pLDDT maps (**b,d**) of psHaloTag1a (**a,b**) and psHaloTag1b (**c,d**). The HaloTag domain is in grey and sAsLOV2 domain in yellow in **a**,**c**. The amino acids corresponding to mutations introduced during rounds of mutagenesis are depicted as sticks.

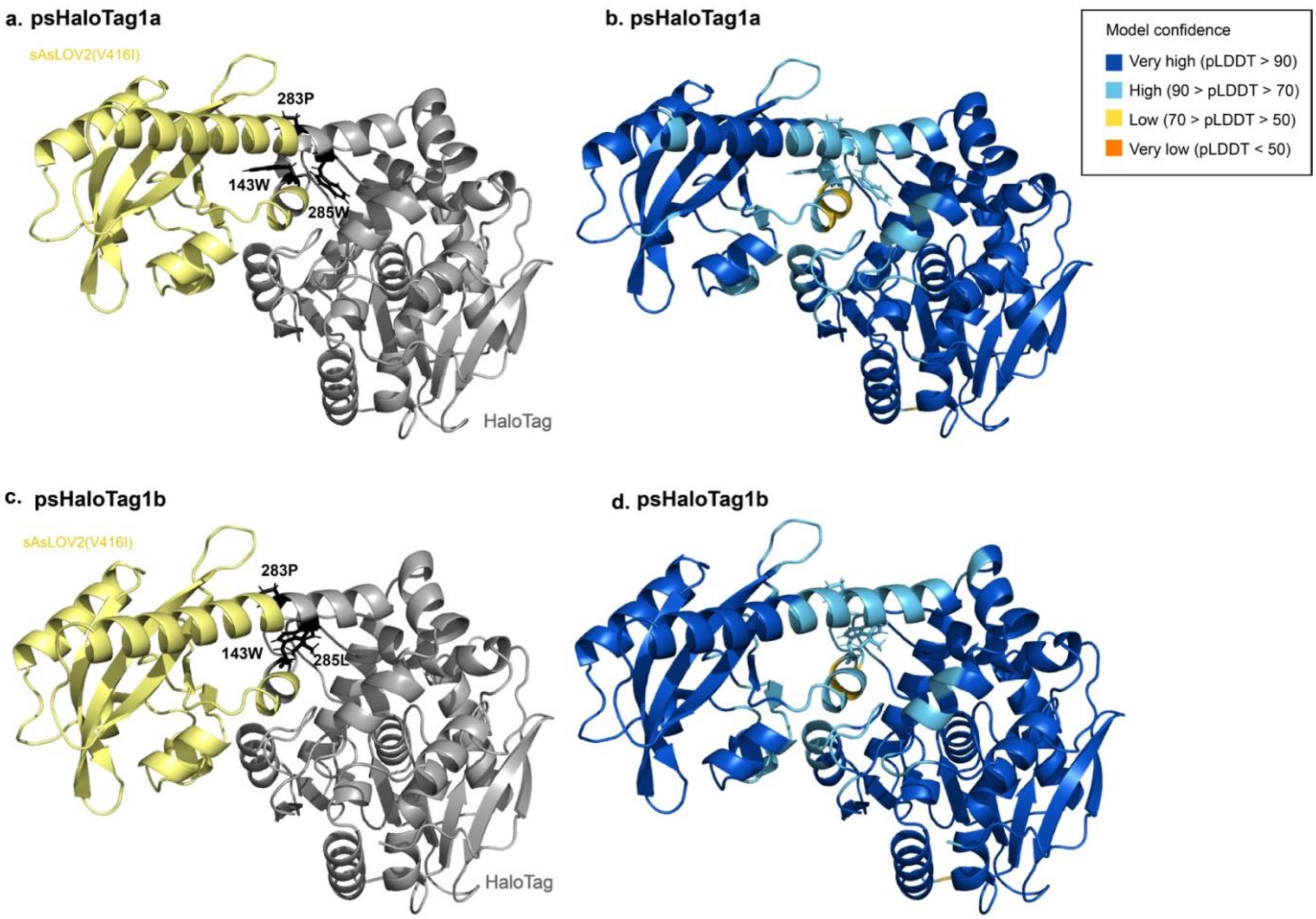

**Figure S12. a.** The crystal structure of psHaloTag1a with **JF$_{635}$-HTL** and FMN contains two chains with one molecule of each psHaloTag1a, **JF$_{635}$-HTL** (magenta), FMN (orange), and $Cl^-$ (cyan) in its asymmetric unit. The two rhodamines (magenta) from chain A (green) and chain B (salmon) are facing each other in the center. **b.** Superimposition of the two assemblies reveals a near identical structure for both protein chains and positioning of the ligands with a root-mean-square deviation (RMSD) of 0.33 Å (for 431 Cα residues). **c.** Comparison of an AlphaFold model (blue) with chain A (green) shows similar folds for the HaloTag and LOV domain part with an RMSD of 1.64 Å (for 371 Cα residues). Here, the HaloTag domain was used for superimposition. A slightly different conformation in the prediction of the Jα helix and the linker region before the Jα helix leads to an offset between the predicted and experimentally determined position of the LOV domains. **d.** Fitting of both FMN molecules into the electron density maps (grey mesh) and the potential covalent binding partner Cys189. Based on the electron density, both FMN molecules do not seem covalently bound. **e.** Fitting of both **JF$_{635}$-HTL** dyes with the released $Cl^-$ and the covalently linked Asp106 residues into the electron density maps (grey mesh). Based on the electron density, the open conformation of the dye fitted best in both chains. The 2m$F_{obs}$-$DF_{calc}$ map is at a contour level of 1 σ.

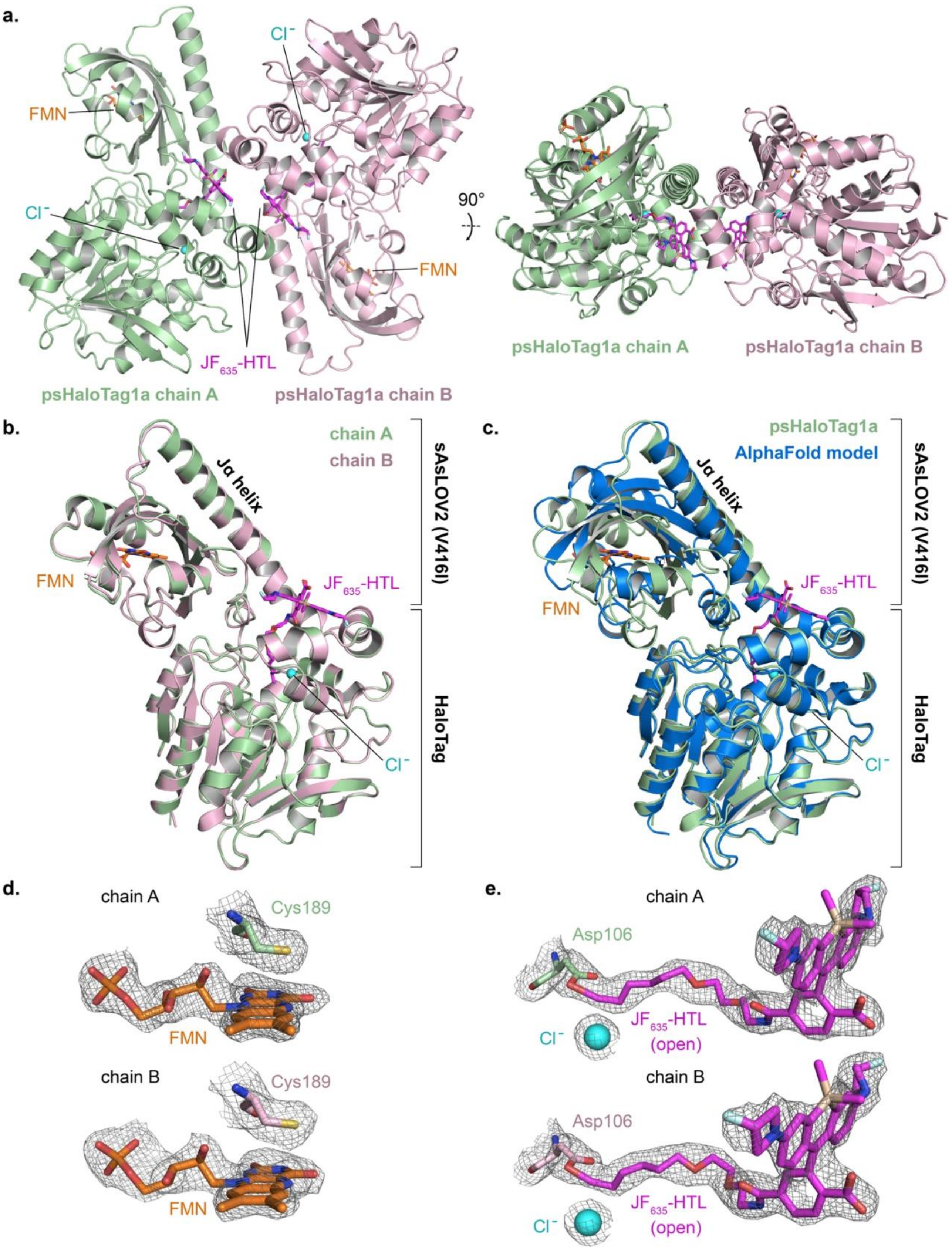
a.
Cl⁻
FMN
Cl⁻
90°
FMN
JF635-HTL
psHaloTag1a chain A
psHaloTag1a chain B
psHaloTag1a chain A
psHaloTag1a chain B
b.
chain A
chain B
Jα helix
sAsLOV2 (V416I)
FMN
JF635-HTL
HaloTag
Cl⁻
c.
psHaloTag1a
AlphaFold model
Jα helix
sAsLOV2 (V416I)
JF635-HTL
FMN
HaloTag
Cl⁻
d.
chain A
Cys189
FMN
chain B
Cys189
FMN
e.
chain A
Asp106
Cl⁻
JF635-HTL
(open)
chain B
Asp106
Cl⁻
JF635-HTL
(open)

**Figure S13. a.** The crystal structure of psHaloTag1a with **JF$_{635}$-HTL** and FMN compares well with available crystal structures of HaloTag7 bound to **JF$_{669}$-HTL** (sand, PDB: 8SW8)[7] and the dark state of AsLOV2 domain (brown, PDB: 2V1A).[8] Superimposition resulted in root-mean-square deviations (RMSD) of 0.50 Å (for 293 Cα residues) for the HaloTag moiety and 1.04 Å (for 137 Cα residues) for the LOV domain, respectively. **b.** Side-view of the superimposed psHaloTag1a structure (green) with HaloTag7 (sand) bound to **JF$_{669}$-HTL**. The position of the rhodamine moieties of **JF$_{635}$-HTL** (magenta) and **JF$_{669}$-HTL** (grey) is not identical, highlighting flexibility. However, as the dyes are surface-exposed, different crystal contacts could affect the different positions. **c.** Electrostatic surface potential representation of psHaloTag1a with the same view as in (b.) displays a negative surface potential around the surface-exposed rhodamine moiety of **JF$_{635}$-HTL**. Electrostatics are displayed in a gradient from negative (red, -5 kT/e) to positive (blue, +5 kT/e) surface potential. **d.** Electrostatic surface potential representation of HaloTag7 with bound **JF$_{669}$-HTL** with the same view as in (b.) displays a negative surface potential around the surface-exposed rhodamine moiety of **JF$_{669}$-HTL**. There is more surface exposure compared to psHaloTag1a (c.) because of the non-present LOV domain. Electrostatics are displayed in a gradient from negative (red, -5 kT/e) to positive (blue, +5 kT/e) surface potential.

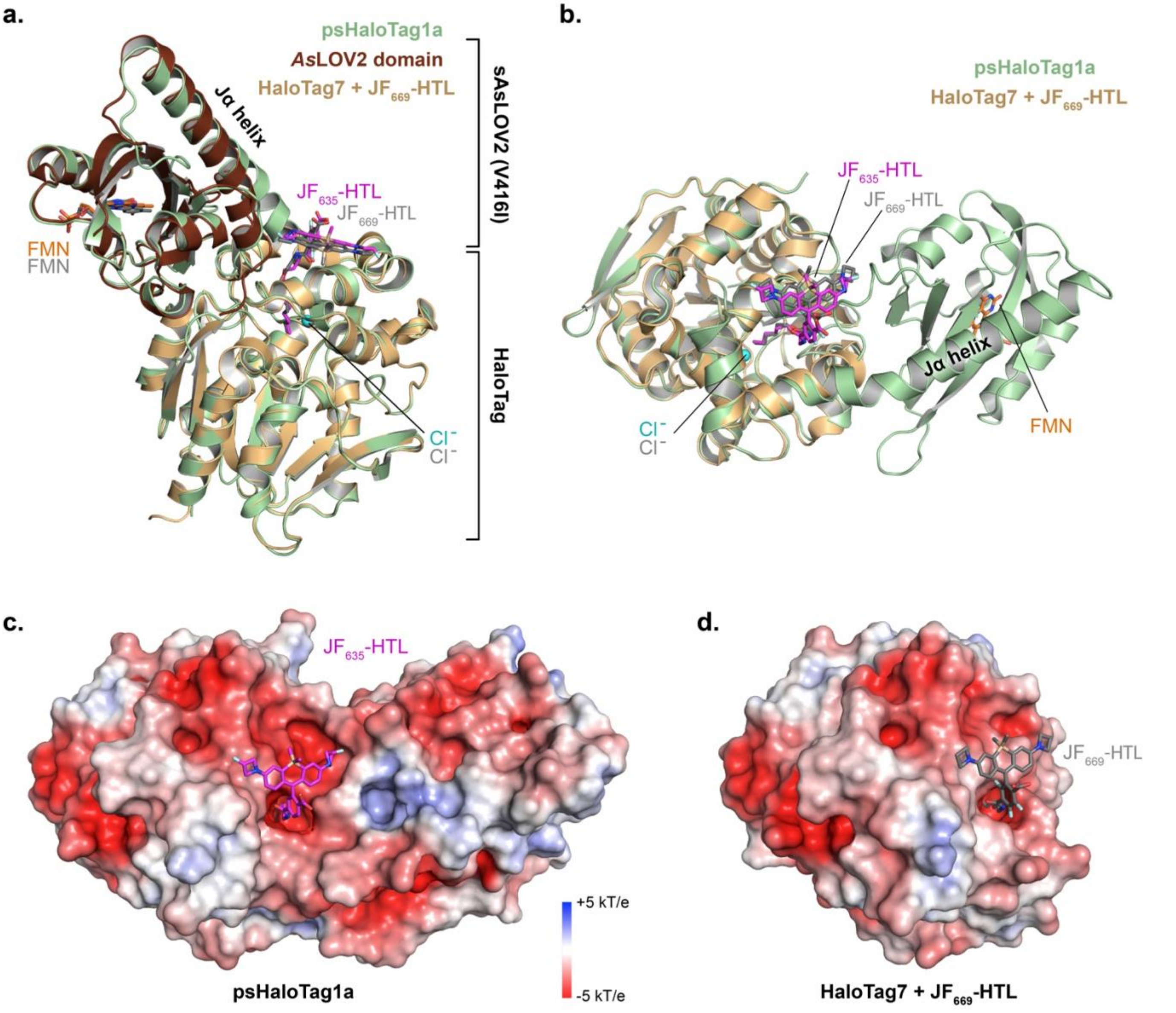

**Figure S14. a.** Representative images of U2OS cells expressing LifeAct-psHaloTag1a-T2A-EGFP and labelled with **JF$_{630}$-HTL**, in the dark (left) and after illumination at 450 nm for 5 minutes (right). Scale bars: 50 μm. **b**. Intensity line profiles corresponding to the images in **a**.

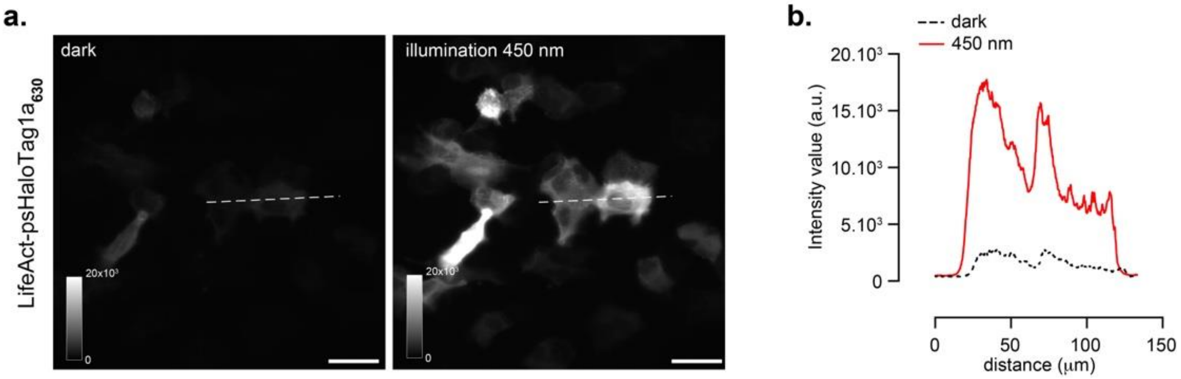

**Figure S15**. SDS-PAGE gel of purified HaloTag (36 kDa) and psHaloTag (52 kDa) proteins.

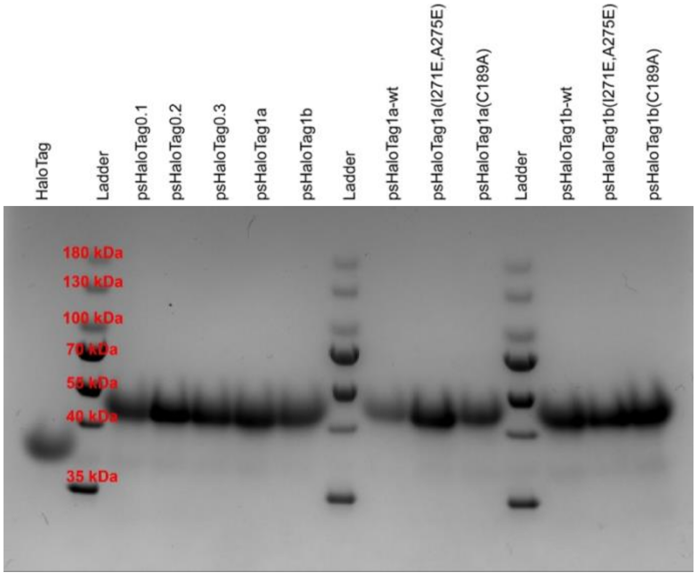

### Supplementary Tables

**Table S1**. Amino acid sequences of psHaloTag constructs. Black = HaloTag, yellow = sAsLOV2, green = inserted amino acid, red = mutation, blue = linker

| |
|---|
| **HaloTag7**:<br>GIGTGFPFDPHYVEVLGERMHYVDVGPRDGTPVLFLHGNPTSSYVWRNIIPHVAPTHRCIAPDLIGMGKSDKPDLGYFFDDHVRFMDAFIEALG<br>LEEVVLVIHDWGSALGFHWAKRNPERVKGIAFMEFIRPIPTWDEWPEFARETFQAFRTTDVGRKLIIDQNVFIEGTLPMGVVRPLTEVEMDHYR<br>EPFLNPVDREPLWRFPNELPIAGEPANIVALVEEYMDWLHQSPVPKLLFWGTPGVLIPPAEAARLAKSLPNCKAVDIGPGLNLLQEDNPDLIGSEI<br>ARWLSTLEI |
| **psHaloTag0.1:**<br>GIGTGFPFDPHYVEVLGERMHYVDVGPRDGTPVLFLHGNPTSSYVWRNIIPHVAPTHRCIAPDLIGMGKSDKPDLGYFFDDHVRFMDAFIEALG<br>LEEVVLVIHDWGSALGFHWAKRNPERVKGIAFMEFIRPIPTWDEWPEGLERIEKNFIITDPRLPDNPIIFASDSFLQLTEYSREEILGRNCRFLQGP<br>ETDRATVRKIRDAIDNQTEVTVQLINYTKSGKKFWNLFHLQPMRDQKGDVQYFIGVQLDGTEHVRDAAEREGVMLIKKTAENIDEAAGFARETF<br>QAFRTTDVGRKLIIDQNVFIEGTLPMGVVRPLTEVEMDHYREPFLNPVDREPLWRFPNELPIAGEPANIVALVEEYMDWLHQSPVPKLLFWGTP<br>GVLIPPAEAARLAKSLPNCKAVDIGPGLNLLQEDNPDLIGSEIARWLSTLEI |
| **psHaloTag0.2:**<br>GIGTGFPFDPHYVEVLGERMHYVDVGPRDGTPVLFLHGNPTSSYVWRNIIPHVAPTHRCIAPDLIGMGKSDKPDLGYFFDDHVRFMDAFIEALG<br>LEEVVLVIHDWGSALGFHWAKRNPERVKGIAFMEFIRPIPTWDEWPEFAGLERIEKNFIITDPRLPDNPIIFASDSFLQLTEYSREEILGRNCRFLQ<br>GPETDRATVRKIRDAIDNQTEVTVQLINYTKSGKKFWNLFHLQPMRDQKGDVQYFIGVQLDGTEHVRDAAEREGVMLIKKTAENIDEAAPFARE<br>TFQAFRTTDVGRKLIIDQNVFIEGTLPMGVVRPLTEVEMDHYREPFLNPVDREPLWRFPNELPIAGEPANIVALVEEYMDWLHQSPVPKLLFWG<br>TPGVLIPPAEAARLAKSLPNCKAVDIGPGLNLLQEDNPDLIGSEIARWLSTLEI |
| **psHaloTag0.3**:<br>GIGTGFPFDPHYVEVLGERMHYVDVGPRDGTPVLFLHGNPTSSYVWRNIIPHVAPTHRCIAPDLIGMGKSDKPDLGYFFDDHVRFMDAFIEALG<br>LEEVVLVIHDWGSALGFHWAKRNPERVKGIAFMEFIRPIPTWDEWPWFAGLERIEKNFIITDPRLPDNPIIFASDSFLQLTEYSREEILGRNCRFLQ<br>GPETDRATVRKIRDAIDNQTEVTVQLINYTKSGKKFWNLFHLQPMRDQKGDVQYFIGVQLDGTEHVRDAAEREGVMLIKKTAENIDEAAPFARE<br>TFQAFRTTDVGRKLIIDQNVFIEGTLPMGVVRPLTEVEMDHYREPFLNPVDREPLWRFPNELPIAGEPANIVALVEEYMDWLHQSPVPKLLFWG<br>TPGVLIPPAEAARLAKSLPNCKAVDIGPGLNLLQEDNPDLIGSEIARWLSTLEI |
| **psHaloTag1a**:<br>GIGTGFPFDPHYVEVLGERMHYVDVGPRDGTPVLFLHGNPTSSYVWRNIIPHVAPTHRCIAPDLIGMGKSDKPDLGYFFDDHVRFMDAFIEALG<br>LEEVVLVIHDWGSALGFHWAKRNPERVKGIAFMEFIRPIPTWDEWPWFAGLERIEKNFIITDPRLPDNPIIFASDSFLQLTEYSREEILGRNCRFLQ<br>GPETDRATVRKIRDAIDNQTEVTVQLINYTKSGKKFWNLFHLQPMRDQKGDVQYFIGVQLDGTEHVRDAAEREGVMLIKKTAENIDEAAPFWRE<br>TFQAFRTTDVGRKLIIDQNVFIEGTLPMGVVRPLTEVEMDHYREPFLNPVDREPLWRFPNELPIAGEPANIVALVEEYMDWLHQSPVPKLLFWG<br>TPGVLIPPAEAARLAKSLPNCKAVDIGPGLNLLQEDNPDLIGSEIARWLSTLEI |
| **psHaloTag1a-wt:**<br>GIGTGFPFDPHYVEVLGERMHYVDVGPRDGTPVLFLHGNPTSSYVWRNIIPHVAPTHRCIAPDLIGMGKSDKPDLGYFFDDHVRFMDAFIEALG<br>LEEVVLVIHDWGSALGFHWAKRNPERVKGIAFMEFIRPIPTWDEWPWFAGLERIEKNFVITDPRLPDNPIIFASDSFLQLTEYSREEILGRNCRFL<br>QGPETDRATVRKIRDAIDNQTEVTVQLINYTKSGKKFWNLFHLQPMRDQKGDVQYFIGVQLDGTEHVRDAAEREGVMLIKKTAENIDEAAPFWR<br>ETFQAFRTTDVGRKLIIDQNVFIEGTLPMGVVRPLTEVEMDHYREPFLNPVDREPLWRFPNELPIAGEPANIVALVEEYMDWLHQSPVPKLLFW<br>GTPGVLIPPAEAARLAKSLPNCKAVDIGPGLNLLQEDNPDLIGSEIARWLSTLEI |
| **psHaloTag1b:**<br>GIGTGFPFDPHYVEVLGERMHYVDVGPRDGTPVLFLHGNPTSSYVWRNIIPHVAPTHRCIAPDLIGMGKSDKPDLGYFFDDHVRFMDAFIEALG<br>LEEVVLVIHDWGSALGFHWAKRNPERVKGIAFMEFIRPIPTWDEWPWFAGLERIEKNFIITDPRLPDNPIIFASDSFLQLTEYSREEILGRNCRFLQ<br>GPETDRATVRKIRDAIDNQTEVTVQLINYTKSGKKFWNLFHLQPMRDQKGDVQYFIGVQLDGTEHVRDAAEREGVMLIKKTAENIDEAAPFLRE<br>TFQAFRTTDVGRKLIIDQNVFIEGTLPMGVVRPLTEVEMDHYREPFLNPVDREPLWRFPNELPIAGEPANIVALVEEYMDWLHQSPVPKLLFWG<br>TPGVLIPPAEAARLAKSLPNCKAVDIGPGLNLLQEDNPDLIGSEIARWLSTLEI |

**psHaloTag1b-wt:**
GIGTGFPFDPHYVEVLGERMHYVDVGPRDGTPVLFLHGNPTSSYVWRNIIPHVAPTHRCIAPDLIGMGKSDKPDLGYFFDDHVRFMDAFIEALG
LEEVVLVIHDWGSALGFHWAKRNPERVKGIAFMEFIRPIPTWDEWPWFAGLERIEKNFVITDPRLPDNPIIFASDSFLQLTEYSREEILGRNCRFL
QGPETDRATVRKIRDAIDNQTEVTVQLINYTKSGKKFWNLFHLQPMRDQKGDVQYFIGVQLDGTEHVRDAAEREGVMLIKKTAENIDEAAPFLR
ETFQAFRTTDVGRKLIIDQNVFIEGTLPMGVVRPLTEVEMDHYREPFLNPVDREPLWRFPNELPIAGEPANIVALVEEYMDWLHQSPVPKLLFW
GTPGVLIPPAEAARLAKSLPNCKAVDIGPGLNLLQEDNPDLIGSEIARWLSTLEI

**psHaloTag1a(C189A):**
GIGTGFPFDPHYVEVLGERMHYVDVGPRDGTPVLFLHGNPTSSYVWRNIIPHVAPTHRCIAPDLIGMGKSDKPDLGYFFDDHVRFMDAFIEALG
LEEVVLVIHDWGSALGFHWAKRNPERVKGIAFMEFIRPIPTWDEWPWFAGLERIEKNFIITDPRLPDNPIIFASDSFLQLTEYSREEILGRNARFLQ
GPETDRATVRKIRDAIDNQTEVTVQLINYTKSGKKFWNLFHLQPMRDQKGDVQYFIGVQLDGTEHVRDAAEREGVMLIKKTAENIDEAAPFWRE
TFQAFRTTDVGRKLIIDQNVFIEGTLPMGVVRPLTEVEMDHYREPFLNPVDREPLWRFPNELPIAGEPANIVALVEEYMDWLHQSPVPKLLFWG
TPGVLIPPAEAARLAKSLPNCKAVDIGPGLNLLQEDNPDLIGSEIARWLSTLEI

**psHaloTag1a(I271E, A275E):**
GIGTGFPFDPHYVEVLGERMHYVDVGPRDGTPVLFLHGNPTSSYVWRNIIPHVAPTHRCIAPDLIGMGKSDKPDLGYFFDDHVRFMDAFIEALG
LEEVVLVIHDWGSALGFHWAKRNPERVKGIAFMEFIRPIPTWDEWPWFAGLERIEKNFIITDPRLPDNPIIFASDSFLQLTEYSREEILGRNCRFLQ
GPETDRATVRKIRDAIDNQTEVTVQLINYTKSGKKFWNLFHLQPMRDQKGDVQYFIGVQLDGTEHVRDAAEREGVMLEKKTEENIDEAAPFWR
ETFQAFRTTDVGRKLIIDQNVFIEGTLPMGVVRPLTEVEMDHYREPFLNPVDREPLWRFPNELPIAGEPANIVALVEEYMDWLHQSPVPKLLFW
GTPGVLIPPAEAARLAKSLPNCKAVDIGPGLNLLQEDNPDLIGSEIARWLSTLEI

**psHaloTag1b(C189A):**
GIGTGFPFDPHYVEVLGERMHYVDVGPRDGTPVLFLHGNPTSSYVWRNIIPHVAPTHRCIAPDLIGMGKSDKPDLGYFFDDHVRFMDAFIEALG
LEEVVLVIHDWGSALGFHWAKRNPERVKGIAFMEFIRPIPTWDEWPWFAGLERIEKNFIITDPRLPDNPIIFASDSFLQLTEYSREEILGRNARFLQ
GPETDRATVRKIRDAIDNQTEVTVQLINYTKSGKKFWNLFHLQPMRDQKGDVQYFIGVQLDGTEHVRDAAEREGVMLIKKTAENIDEAAPFLRE
TFQAFRTTDVGRKLIIDQNVFIEGTLPMGVVRPLTEVEMDHYREPFLNPVDREPLWRFPNELPIAGEPANIVALVEEYMDWLHQSPVPKLLFWG
TPGVLIPPAEAARLAKSLPNCKAVDIGPGLNLLQEDNPDLIGSEIARWLSTLEI

**psHaloTag1b(I271E, A275E):**
GIGTGFPFDPHYVEVLGERMHYVDVGPRDGTPVLFLHGNPTSSYVWRNIIPHVAPTHRCIAPDLIGMGKSDKPDLGYFFDDHVRFMDAFIEALG
LEEVVLVIHDWGSALGFHWAKRNPERVKGIAFMEFIRPIPTWDEWPWFAGLERIEKNFIITDPRLPDNPIIFASDSFLQLTEYSREEILGRNCRFLQ
GPETDRATVRKIRDAIDNQTEVTVQLINYTKSGKKFWNLFHLQPMRDQKGDVQYFIGVQLDGTEHVRDAAEREGVMLEKKTEENIDEAAPFLRE
TFQAFRTTDVGRKLIIDQNVFIEGTLPMGVVRPLTEVEMDHYREPFLNPVDREPLWRFPNELPIAGEPANIVALVEEYMDWLHQSPVPKLLFWG
TPGVLIPPAEAARLAKSLPNCKAVDIGPGLNLLQEDNPDLIGSEIARWLSTLEI

**Table S2**. Apparent kinetic rate constants of the photoswitching (upon illumination at 450 nm) and thermal relaxation (in the dark at room temperature) of psHaloTag variants labelled with **$JF_{635}$-HTL**, for both the FMN cofactor signal and **$JF_{635}$** signal. The corresponding full kinetic profiles can be found in Figure S5. n.m.: not measured.

| | Illumination at 450 nm | | Thermal relaxation (dark, room temperature) | | | |
|---|---|---|---|---|---|---|
| Construct | $k_{ON}(JF_{635})$ ($min^{-1}$) | $t_{1/2, ON}(JF_{635})$ (min) | $k_{relax}$(FMN) ($min^{-1}$) | $t_{1/2,relax}$(FMN) (min) | $k_{relax}(JF_{635})$ ($min^{-1}$) | $t_{1/2,relax}(JF_{635})$ (min) |
| psHaloTag0.1 | n.m. | n.m. | 0.091 | 7.6 | 0.103 | 6.8 |
| psHaloTag0.2 | n.m. | n.m. | 0.088 | 7.9 | n.m. | n.m. |
| psHaloTag0.3 | 0.030 | 24.1 | 0.090 | 7.7 | 0.055 | 12.7 |
| psHaloTag1a | 0.048 | 14.3 | 0.088 | 7.9 | 0.036 | 19.5 |
| psHaloTag1a-wt | 0.039 | 17.0 | 1.20 | 0.6 | 0.117 | 5.9 |
| psHaloTag1b | 0.307 | 2.3 | 0.104 | 6.7 | 0.072 | 9.7 |
| psHaloTag1b-wt | 0.332 | 2.1 | 1.22 | 0.6 | 0.378 | 1.8 |

**Table S3.** Crystallographic data and refinement statistics. Data for highest-resolution shell are shown parenthesis.

| | **psHaloTag1a$_{635}$** |
|---|---|
| **Data collection** | |
| Space group | C121 |
| Resolution (Å) | 70.36 – 2.4 (2.49 – 2.4) |
| Unique reflection | 40131 (3989) |
| a, b, c (Å) | 167.74, 78.02, 81.81 |
| α, β, γ (°) | 90.0, 103.9, 90.0 |
| $R_{merge}$ | 0.262 (1.228) |
| $R_{pim}$ | 0.106 (0.491) |
| Mean (I/σ(I)) | 5.6 (1.04) |
| Multiplicity | 7.0 (7.1) |
| Completeness (%) | 99.55 (99.60) |
| $CC_{1/2}$ | 0.99 (0.58) |
| **Refinement** | |
| $R_{work}$ (%) | 21.31 (30.16) |
| $R_{free}$ (%) | 25.23 (32.81) |
| RMSD Bond length (Å) | 0.009 |
| RMSD Bond angle (°) | 1.35 |
| Ramachandran favored (%) | 95.35 |
| Ramachandran allowed (%) | 4.65 |
| Ramachandran outliers (%) | 0.0 |
| Rotamer outliers (%) | 2.26 |
| Clashscore | 7.01 |
| Average B factor (Å$^2$) | 39.91 |
| Protein | 39.75 |
| Ligands | 43.83 |
| Solvent | 40.50 |
| PDB identifier | 9HKF |

**Table S4.** List and composition of buffers used in this work.

| Buffer | Composition |
|---|---|
| Spectroscopy buffer | 20 mM Tris-HCl pH 7.4, 100 mM NaCl, 0.1 mg·ml$^{-1}$ CHAPS |
| Lysis buffer | 20 mM Tris-HCl pH 7.4, 300 mM NaCl, 1 mM PMSF, 0.01 mg·mL$^{-1}$ DNase [for screening assays 2 mg·mL$^{-1}$ lysozyme/for protein purification ½ tablet of protease inhibitor (cOmplete)] |
| Ni-NTA chromatography wash buffer | 20 mM Tris HCl pH 7.4, 300 mM NaCl, 10 mM imidazole |
| Ni-NTA chromatography elution buffer | 20 mM Tris-HCl pH 7.4, 300 mM NaCl, 500 mM imidazole (150 mM imidazole for protein crystallization) |
| Dialysis buffer | 20 mM Tris-HCl pH 7.4, 100 mM NaCl, 5 mM imidazole, 2 mM β-mercaptoethanol |
| Size exclusion chromatography (SEC) buffer/Storage buffer | 20 mM Tris-HCl pH 7.4, 100 mM NaCl |
| DMEM | DMEM high glucose (4.5 g·L$^{-1}$), 10% fetal bovine serum, 2 mM L-glutamine, 1 mM sodium pyruvate, 100 U·mL$^{-1}$ penicillin, 100 µg·mL$^{-1}$ streptomycin |
| Imaging buffer (microscopy) | DMEM without phenol red high glucose (4.5 g·L$^{-1}$), 10% fetal bovine serum, 1 mM sodium pyruvate, 1 mM L-glutamine |

**Table S5.** Thermal stability of purified proteins determined by differential scanning fluorimetry.

| Protein variant | $T_{onset}$ (°C) | $T_m$ (°C) | $T_{agg}$ (°C) |
|---|---|---|---|
| psHaloTag0.1 | 34.8 | 46.4 | 42.5 |
| psHaloTag0.2 | 37.9 | 48.5 | 41.2 |
| psHaloTag0.3 | 35.3 | 46.3 | 40.6 |
| psHaloTag1a | 40.8 | 48.6 | 39.7 |
| psHaloTag1b | 37.6 | 48.6 | 40.7 |
| psHaloTag1a-wt | 42.2 | 49.8 | 41.9 |
| psHaloTag1b-wt | 39.1 | 48.7 | 40.7 |
| psHaloTag1a (I271E, A275E) | 35.4 | 45.2 | 40.4 |
| psHaloTag1a (C189A) | 41.4 | 49.0 | 40.2 |
| psHaloTag1b (I271E, A275E) | 33.9 | 45.0 | 44.3 |
| psHaloTag1b (C189A) | 37.9 | 47.6 | 42.4 |

**Table S6**. Plasmids used for cell culture and microscopy.

| Cellular target | Construct | Addgene # |
|---|---|---|
| Nucleus | pCDNA5_FRT_TO_H2B_HaloTag_T2A_EGFP | 135444 |
| | pCDNA5_FRT_TO_H2B_psHaloTag1a_T2A_EGFP | - |
| Mitochondria | pCDNA5_FRT_TO_TOMM20_HaloTag_T2A_EGFP | 135443 |
| | pCDNA5_FRT_TO_TOMM20_ psHaloTag1a _T2A_EGFP | - |
| Actin cytoskeleton | pCDNA5_FRT_TO_LifeAct_ HaloTag _T2A_EGFP | 135445 |
| | pCDNA5_FRT_TO_LifeAct_ psHaloTag1a _T2A_EGFP | - |

### General Experimental Information

#### Chemicals, reagents and buffers

**$JF_{635}$-HTL** was synthesized as previously described.[2] The HaloTag ligands of **$JF_{525}$**,[2] **$JF_{526}$**,[3] **$JF_{585}$**,[2] **$JF_{626}$**,[4] **$JF_{629}$**,[4] **$JF_{630}$**,[4] **$JF_{639}$**,[4] **$JF_{711}$**[5] were provided by the Lavis group (Janelia Research Campus, HHMI). The HaloTag ligands of **$MaP_{618}$**[6] and **$MaP_{700}$**[6] were provided by the Johnsson group (Max Planck Institute for Medical Research).

Commercial reagents were obtained from reputable suppliers and used as received.
The composition of buffers used in this work can be found in Table S4.

### UV-Vis and Fluorescence Spectroscopy

All measurements were performed at room temperature (23 ± 2°C). Fluorophore-ligands were initially prepared as 1 mM stock solutions in DMSO (spectroscopy grade) and subsequently diluted in appropriate solvents and buffers to ensure that the final DMSO concentration did not exceed 1% v/v. Unless otherwise specified, spectroscopic measurements with purified proteins were performed in aqueous buffer containing 20 mM Tris-HCl and 100 mM NaCl at pH 7.4, to which 0.1 mg·mL$^{-1}$ 3-((3-cholamidopropyl)dimethylammonio)-1-propanesulfonate (CHAPS) was added. Spectroscopic measurements were conducted using 1-cm path-length quartz cuvettes (Hellma Suprasil Quartz), or in 96-well plates (Nunc™ MicroWell™ 96-Well, Nunclon Delta-Treated, Flat-Bottom Microplate).

Absorption spectra were recorded using a Cary Model 60 spectrophotometer (Agilent). Absorption spectra, including maximum absorption wavelength ($\lambda_{max}$), extinction coefficient (ε), and maximum emission wavelength ($\lambda_{em}$), were measured in triplicates. Spectra were recorded at 5 µM dye concentration, unless otherwise stated. For measurements in the presence of proteins, the dye ligand was incubated with 1.5 equivalents (eq) of purified protein for at least 3 h in the dark at room temperature.

Fluorescence spectra were recorded using a JASCO spectrofluorometer (FP-8500), at 1 µM dye concentration of and 1.5 µM protein concentration, unless otherwise stated. Fluorescence intensity measurements at a single wavelength were performed with a TECAN plate reader (TECAN Infinite M1000 Pro fluorescence plate reader or TECAN Spark microplate reader) equipped with appropriate filters and a monochromator.

Data analysis and graph plotting were carried out using Prism (GraphPad).

The following notations are used:

| | |
|---|---|
| $\lambda_{max}$ (nm) | Wavelength at maximal absorption |
| $\lambda_{em}$ (nm) | Wavelength at maximal fluorescence emission |
| ε ($M^{-1}\cdot cm^{-1}$) | Molar extinction coefficient |
| $\Phi_F$ | Fluorescence quantum yield |
| $A/A_0$ | Change in absorbance |
| $F/F_0$ | Change in fluorescence |
| $k_{ON}$ ($min^{-1}$) | Apparent rate constant of the turn-on photoinduced process |

$k_{relax}$ ($min^{-1}$) Apparent rate constant of thermal relaxation
$t_{1/2,ON}$ (min) Half-life of the turn-on, photoinduced process. Determined as $t_{1/2,ON} = \ln 2/ k_{ON}$.
$t_{1/2, relax}$ (min) Half-life of the thermal relaxation. Determined as $t_{1/2, relax} = \ln 2/k_{relax}$.

#### Extinction coefficient measurements

Purified protein solutions were labelled with 0.7 eq **JF$_{635}$-HTL** in aqueous buffer (20 mM Tris-HCl, 100 mM NaCl, 0.1 mg·mL$^{-1}$ CHAPS, pH 7.4) for 3 h at room temperature, in the dark. The absorption spectra of labelled protein solutions at different concentrations (1 µM, 1.25 µM, 1.5 µM, 2.0 µM, 2.5 µM, 3 µM, 4 µM, 5 µM, and 6 µM) were recorded before and after illumination at 450 nm. The spectra were baseline corrected, and the maximum absorption values were plotted against concentration. The extinction coefficient (ε) at $\lambda_{max}$ was determined by linear curve fitting using the Beer-Lambert law.

#### Quantum yield measurements

Fluorescence quantum yields ($\Phi_F$) were measured using an absolute quantum yield measurement system (Quantaurus, C11347, Hamamatsu). Measurements were carried out using dilute samples (A < 0.1). Self-absorption corrections were performed using the Quantaurus software. Purified protein solutions (1.5 µM) were labelled with **JF$_{635}$-HTL** (1 µM) in aqueous buffer (20 mM Tris-HCl, 100 mM NaCl, 0.1 mg·mL$^{-1}$ CHAPS, pH 7.4) for 3 h at room temperature in the dark before measurements. Afterward, the samples were illuminated at 450 nm until complete photoswitching was achieved (assessed by UV-Vis spectroscopy), and the quantum yield of the ON state was measured.

#### Photoswitching

Photoswitching was performed using a LED illumination device, designed and built by the Electronical Workshop at EMBL Heidelberg (Figure S3). They consist of two parts: the control board and the LED board, comprised of 24 LEDs at 450 nm (LED450-03, Roithner LaserTechnik). The illumination device could be controlled using the RealTerm Serial terminal program, with the following parameters adjustable: power intensity, exposure time ON and OFF (in ms), and number of ON/OFF cycles. For all experiments, the maximum power intensity of the system was used, corresponding to 2.6 mW.cm$^{-2}$ in our setup.

#### Thermal relaxation kinetics

Thermal relaxation kinetics measurements were performed in a microplate reader (TECAN Infinite M1000 microplate reader) at room temperature, in the dark, in 96-well plates, in aqueous buffer (20 mM Tris-HCl, 100 mM NaCl, 0.1 mg·mL$^{-1}$ CHAPS, pH 7.4). The plate was illuminated at 450 nm until the maximal ON state has been reached (assessed by fluorescence measurement). The fluorescence intensity values of the fluorophore ($\lambda_{max}/\lambda_{em}$ = 640/660 nm) and the FMN ($\lambda_{max}/\lambda_{em}$ = 450/525 nm) were recorded every minute for a duration of 2 h. The resulting experimental data were fitted to a one-phase exponential decay function to determine the FMN fluorescence increase rate and the dye fluorescence decrease rate as a proxy for the FMN-Cys adduct lifetime and the conformational relaxation of the HaloTag binding cavity. The following equation was used:

$$F(t) = (F_{MAX} - F_0) + exp(-k_{relax} * t) + F_0$$

Where:

t is the time in min.
F(t) is the fluorescence intensity of the sample at t.

$F_{MAX}$ is the fluorescence intensity at t = 0 (illuminated state).
$F_0$ is the fluorescence intensity in the dark state.
$k_{relax}$ is the apparent rate constant of the thermal relaxation, in $min^{-1}$.

#### pH stability

Aqueous buffers of varying pH (2-11) were prepared using $Na_2HPO_4$ (200 mM) and sodium citrate (100 mM) for pH 2-7 and $Na_2CO_3$ (100 mM) and $NaHCO_3$ (100 mM) for pH 8-11, respectively. Purified psHaloTag 1a prelabelled with **$JF_{635}$-HTL** (1 µM) was incubated with the respective pH-adjusted buffer for 3 h and the emissions of FMN ($\lambda_{exc}/\lambda_{em}$ = 450/525 nm) and **$JF_{635}$** ($\lambda_{exc}/\lambda_{em}$ = 640/660 nm) before and after illumination (50 min, 450 nm, 2.6 mW·cm$^{-2}$) were measured on a microplate reader (Tecan Spark) before and after illumination (50 min, 450 nm, 2.6 mW.cm$^{-2}$). Following illumination, the thermal relaxation was determined by recording the emission of FMN and **$JF_{635}$** in time intervals of 5 min for a total duration of 90 min. The thermal relaxation constants were determined as described above.

#### Binding kinetics

Binding kinetics measurements were performed in Cary Model 60 spectrophotometer (Agilent) at room temperature, in the dark, 1-cm path-length quartz cuvettes (Hellma Suprasil Quartz), in aqueous buffer (20 mM Tris-HCl, 100 mM NaCl, 0.1 mg·mL$^{-1}$ CHAPS, pH 7.4). To **$JF_{635}$-HTL** (5 µM) in aqueous buffer was psHaloTag1a (7.5 µM) added in the dark and quickly mixed. The absorption spectra were recorded every minute for a duration of 60 minutes. The resulting experimental data were fitted to a mono exponential function. The following equation was used:

$$F(t) = (F_{MAX} - F_0) * (1 - exp(-k_{app} * t)) + F_0$$

Where:
t is the time in min.
F(t) is the absorption intensity of the sample at t.
$F_{MAX}$ is the absorption intensity in the fully bound state
$F_0$ is the absorption intensity prior binding
$k_{app}$ is the apparent rate constant of binding, in $min^{-1}$.

### Cloning, Screening, Protein Expression and Purification

#### Cloning

The pET51b(+) vector (pET51b(+)-HaloTag7, Addgene #167266) was kindly provided by K. Johnsson (Max Planck Institute for Medical Research) and was used for cloning and protein expression in *E. coli* BL21(DE3). Proteins were N-terminally tagged with Hisx10, followed by a tobacco etch virus (TEV) protease cleavage site (ENLYFQ|G). For *in vitro* screening, YFP was fused to the C-terminus of psHaloTag constructs. For crystallization purposes, the pCoofy1 vector (Addgene #43974, kindly provided by the Protein Expression and Purification Core Facility at EMBL Heidelberg) was selected. In this case, proteins were N-terminally tagged with Hisx6, followed by the HRV 3C protease cleavage site (LEVLFQ|GP). For mammalian cell experiments, the pcDNA5/FRT/TO vector (ThermoFisher Scientific) containing HaloTag7 fused to different subcellular targeting motifs (H2B, TOMM20, or LifeAct) were used for cloning and imaging (kindly provided by K. Johnsson (Max Planck Institute for Medical Research)).

The amino acid sequences of the isolated psHaloTag variants can be found in Table S1. For cloning, the psHaloTag nucleic acid sequence was amplified by polymerase chain reaction (PCR) and inserted into the desired vector backbone via Gibson assembly (New England Biolabs) and purified by DNA Clean & Concentrator-5 (Zymo Research). In case of point mutations (site-directed or site-saturation mutagenesis for library generation) the desired codon or a NNS degenerate codon was used at the 5´-of the forward primer for construct amplification by PCR. Generated products were then ligated using KLD mix (New England Biolabs) and purified by DNA Clean & Concentrator-5 (Zymo Research). Plasmid DNA was subsequently electroporated in *E. coli* Turbo cells (New England Biolabs) plated on LB agar plates with appropriate antibiotics and incubated at 37°C overnight. In all cases, plasmid DNA was extracted from individual bacterial colonies (QIAwave Plasmid Miniprep Kit, Qiagen), and the sequences of generated plasmids were verified by Sanger sequencing (Eurofins Genomics).

#### Site-saturation mutagenesis (SSM) for library generation

Site-saturation mutagenesis was performed using NNS degenerate codon at the 5´-of the forward primer for construct amplification by PCR. Generated products were then ligated using KLD mix (New England Biolabs) and purified by DNA Clean & Concentrator-5 (Zymo Research). Generated DNA constructs by site-saturation mutagenesis were electroporated into *E. coli* strain BL21(DE3). The cells were plated on agar LB plates bearing 100 µg·mL$^{-1}$ ampicillin and 200 µM isopropyl-β-D-thiogalactopyranoside (IPTG) and incubated at 37°C overnight. Diversity was proven by plasmid extraction from a selective liquid LB culture (in autoinduction medium containing 100 µg·mL$^{-1}$ ampicillin) grown at 37°C overnight. The plasmid libraries were sequenced using the Sanger method (Eurofins Genomics) to verify the proper incorporation of random codons at the desired position. Single colonies were used to inoculate 1 mL selective auto-induction medium containing 100 µg·mL$^{-1}$ ampicillin in a U-bottom-shaped 96-deep well plates (ThermoFisher Scientific). The autoinduction medium was prepared in-house.[9]

Three wells of the 96-well plate were reserved for parental HaloTag7 as a positive control, three wells for the parent psHaloTag construct as a comparison for $F/F_0$, and three wells for sAsLOV2 as a negative control for the binding assay. The bacterial cultures were incubated at room temperature for 48 h with shaking at 350 rpm. Subsequently, 40 µL of the cultures were used to inoculate 40 µL of sterile LB medium containing 30% glycerol in a 384-well plate and kept at -70°C as archives. The rest of the cultures were harvested by centrifugation (3500 rpm, 15 min, 4°C). The bacterial pellets in each well were resuspended in 500 µL lysis buffer (20 mM Tris-HCl, 300 mM NaCl, 2 mg·mL$^{-1}$ lysozyme, 0.01 mg·mL$^{-1}$ DNase) and were submitted to five cycles of freeze/thawing with liquid nitrogen before resuspension and incubating at room temperature overnight at 350 rpm shaking. The obtained cell lysate was cleared by centrifugation (3500 rpm, 20 min, 4°C). The cleared supernatant used for the screening assay.

#### Screening assay

Cleared supernatants from cell lysates (95 µL) were transferred into a 96-well plate (Nunc™ MicroWell™ 96-Well, Nunclon Delta-Treated, Flat-Bottom Microplate). Basal **JF$_{635}$** ($\lambda_{exc}/\lambda_{em}$ = 640/660 nm) and YFP ($\lambda_{exc}/\lambda_{em}$ = 515/530 nm) fluorescence were measured in a microplate reader (TECAN Infinite M1000 microplate reader) before adding **JF$_{635}$-HTL** in each well (5 µL of DMSO stock solution, 50 nM final concentration). Wells with a protein concentration < 100 nM were discarded from the analysis. After incubation at room temperature in the dark for 4 h while shaking, fluorescence intensities were measured to obtain the $F_0$ values. The plate was then illuminated three times for 3 min at 450 nm (LED box, 2.6 mW·cm$^{-2}$), and the fluorescence intensities were measured subsequently after each illumination. To verify and quantify binding of the fluorophore to the protein, a second plate was prepared in the same way but instead of illuminating it, purified HaloTag protein (100 nM final concentration) was added. After 30 min

incubation, the fluorescence intensities were measured again. Efficient ligand binding was confirmed if no increase in fluorescence intensity was observed following addition of HaloTag protein, ensuring that no free ligand was left unbound in the lysate. The control wells were used to determine the mean and SD of YFP and **$JF_{635}$** fluorescence intensities of the parental proteins as well as $F/F_0$ upon illumination. Wells with **$JF_{635}$** fluorescence intensities showing a higher $F/F_0$ upon illumination than the control were selected for further characterization. A representation of the screening pipeline can be found in Figure S2.
Plasmids of selected wells were retrieved from stored bacterial archives and sequenced. Their photoswitching properties were then confirmed, with selected variants expressed and lysed from 50 mL selective LB cultures (as described above). Each protein (1 µM) was labelled with **$JF_{635}$-HTL** (50 nM) in 100 µL buffer in a 96-well plate and incubated for 4 h at room temperature in the dark. The **$JF_{635}$** and YFP fluorescence intensities were measured as previously described. Measurements were performed in triplicates. Mean and 90% confidence intervals were calculated for every variant and compared with those of the parental protein. Variants with significant changes in **$JF_{635}$** fluorescence intensity upon illumination (450 nm, 2.6 $mW \cdot cm^{-2}$) compared with the parental protein were selected for further purification and characterization.

**Protein expression and purification**

Proteins were expressed using the pET51b(+) plasmids in *E. coli* BL21(DE3) cells. LB cultures containing 100 $\mu g \cdot mL^{-1}$ ampicillin were grown at 37°C to an optical density at 600 nm (OD600) of 0.4-0.6, induced by the addition of IPTG (1 mM final concentration) for protein expression and further grown at 18°C overnight (shaking 200 rpm). The cells were harvested by centrifugation (6000 rpm, 20 min, 4°C) and lysed by microfluidizer (Microfluidics Microfluidizer, M-110L FluidProcessor) in lysis buffer (20 mM Tris-HCl pH 7.4, 300 mM NaCl, 1 mM PMSF, ½ tablet of protease inhibitor, 0.01 $mg \cdot mL^{-1}$ DNase). The cell lysate was cleared by ultra-centrifugation (35000 rpm, 45 min, 4°C). Proteins were purified using affinity-tag Ni-NTA agarose (Qiagen) and the eluted fractions (20 mM Tris-HCl pH 7.4, 300 mM NaCl, 500 mM imidazole) were then pooled. Size exclusion chromatography (SEC) (HiLoad 16/600 Superdex 200 pg, Cytiva) was performed to obtain the monomeric fraction with elution using SEC buffer (20 mM Tris-HCl, 100 mM NaCl, pH 7.4). This final protein fraction was concentrated (Amicon Ultra Centrifugal Filter, 30 kDa MWCO, Merck) to 100-170 µM and stored in 100 µL aliquots at −70°C. These proteins were also further characterized by NanoDSF (Prometheus NT.Flex, see below), mass photometry (Refeyn $Two^{MP}$ from Refeyn), and SDS-PAGE gel (Bio-Rad, Mini-PROTEAN® TGX™ stain-free gels, Figure S15).

**Protein stability experiments**

The temperature-dependent unfolding, as a proxy for thermal stability, was determined by using differential scanning fluorimetry (DSF), by employing a Prometheus NT.Flex (NanoTemper Technologies GmbH, Munich, Germany) instrument. DSF monitors the unfolding of proteins by detecting temperature-dependent changes in aromatic amino acid fluorescence. Purified psHaloTag samples (10 µL, 0.4–0.5 $mg \cdot mL^{-1}$) were subjected to a linear temperature increase (20–95°C, $1°C \cdot min^{-1}$). The intrinsic tryptophan fluorescence of the protein was monitored continuously (18 data points per minute) at 330 and 350 nm. Unfolding transition midpoints (expressed as melting temperature, Tm) were determined from the first derivative of the fluorescence ratio (F350/F330) by using the RT.ThermControl Software (NanoTemper Technologies GmbH). This was done by Karine Lapouge from the Protein Expression and Purification Core Facility at EMBL Heidelberg. Values can be found in Table S5.

### Structure Prediction

ColabFold is an accessible tool for predicting protein structures that combines the fast homology search capabilities of MMseqs2 with AlphaFold2 or RoseTTAFold algorithms.[10] ColabFold also gives important confidence measures for evaluating the predicted protein structure: pLDDT (predicted Local Distance Difference Test) and PAE (Predicted Aligned Error). pLDDT scores range from 0 to 100, indicating confidence in the accuracy of individual residues: scores above 90 suggest high confidence, while scores below 70 indicate potential inaccuracies. PAE measures the expected positional error between pairs of residues, with lower values reflecting greater confidence in their spatial arrangement. For 3D protein structure prediction, protein sequences were inputted into ColabFold version 1.5.2 (https://github.com/sokrypton/ColabFold) installed on the local HPC cluster through EasyBuild environment module (https://docs.easybuild.io/version-specific/supported-software/c/ColabFold/) with the help of Federico Marotta (Bork Group, EMBL Heidelberg). For each sequence, five models were generated and ranked by their pLDDT scores. The predicted structures were visualized and analyzed using PyMOL.

### X-Ray crystallography

#### Protein purification and crystallization

pCoofy1-derived vector with kanamycin antibiotic resistance (Addgene #43974, kindly provided by the Protein Expression and Purification Core Facility at EMBL Heidelberg) was used for protein crystallization purposes. In this case, psHaloTag1a was N-terminally tagged with Hisx6, followed by the HRV 3C protease cleavage site (LEVLFQ|GP). psHaloTag1a was expressed in *E. coli* BL21(DE3) cells. 3 liters LB cultures were grown in the presence of 100 µg·mL$^{-1}$ kanamycin at 37°C to an optical density at 600 nm of 0.4−0.6, induced by the addition of IPTG (1 mM final concentration) and further grown at 18°C overnight (200 rpm). The cells were harvested by centrifugation (6000 rpm, 20 min, 4°C) and lysed by microfluidizer (Microfluidics Microfluidizer, M-110L FluidProcessor) in lysis buffer (20 mM Tris-HCl pH 7.4, 300 mM NaCl, 1 mM PMSF, ½ tablet of protease inhibitor (cOmplete, Roche), 0.01 mg·mL$^{-1}$ DNase (Roche)). The cell lysate was cleared by ultra-centrifugation (35000 rpm, 45 min, 4°C). Proteins were purified using Ni-NTA chromatography (Protino Ni-NTA 5 mL FLPC column, Macherey-Nagel) and the eluted fraction (20 mM Tris-HCl pH 7.4, 300 mM NaCl, 150 mM imidazole) was dialyzed overnight in the presence of His-tagged HRV 3C protease (1:200) in dialysis buffer (50 mM Tris-HCl pH 7.4, 100 mM NaCl, 5 mM imidazole, 2 mM β-mercaptoethanol) followed by a reverse Ni-NTA chromatography. The flow-through fraction was then used for size exclusion chromatography. The obtained monomeric fraction was then incubated with **$JF_{635}$-HTL** (in excess, 2 molar equivalents) overnight at 4°C in SEC buffer containing 1% DMSO. Excess dye was then removed by PD-10 desalting column (Cytiva) and dye-labelled protein was concentrated to 47 mg·mL$^{-1}$ for protein crystallization screens. $psHaloTag1a_{635}$ was screened for crystallization using the sitting drop vapor diffusion method with different commercial crystallization screens. Sample and crystallization conditions were mixed in 96-well plates with two sample concentrations (23 mg·mL$^{-1}$ and 12 mg·mL$^{-1}$ final concentrations) by pipetting 100 nL of sample to 100 nL crystallization buffer using a Mosquito LCP pipetting robot (SPT Labtech). Crystallization was carried out at 20°C in a crystallization incubator (Formulatrix). Diffracting crystals grew in a condition with 0.1 M Bis-Tris propane pH 6.5, 0.2 M sodium nitrate, 20% (w/v) PEG3350, 10% (v/v) ethylene glycol (Ligand Friendly Screen, Molecular Dimensions). Crystals appeared after 1-5 days, were fished in CrystalCap SPINE HT loops, and frozen with an additional 20 % (v/v) glycerol as cryo-protectant in liquid nitrogen.

#### X-Ray data collection and model building

psHaloTag1a$_{635}$ X-ray diffraction data set was collected at beamline ID23-2 (ESRF, Grenoble) at 100 K with reflections up to 2.4 Å. The crystal belonged to the C121 space group with unit cell dimensions of 167.7 Å x 78.0 Å x 81.8 Å and 90°, 104°, 90° angles. Reflections on diffraction images were integrated by the ESRF automated processing pipeline with XDS[11] and STARANISO.[12] Subsequently, phases were obtained by molecular replacement using a HaloTag protein model (PDB: 5Y2Y)[13] with PHASER-MR[14] implemented in the PHENIX package.[15] Iterative model building and refinement were performed with Coot[16] and Phenix.refine.[17] The asymmetric unit comprises two molecules of psHaloTag1a$_{635}$.

### Cell Culture and Microscopy

#### Cell culture

U2OS cells (ATCC) were cultured in Dulbecco's modified Eagle medium (DMEM, high glucose (4.5 g·L$^{-1}$) with phenol red (Gibco), supplemented with 10% v/v fetal bovine serum (ThermoFisher Scientific), penicillin (100 units·mL$^{-1}$), streptomycin (100 µg·mL$^{-1}$, Gibco), 2 mM L-Glutamine (Gibco) and 1 mM sodium pyruvate (Gibco), and maintained at 37°C in a humidified 5% v/v $CO_2$ environment.

Plasmids used can be found in Table S6. Transient transfections were performed using FUGENE6 (Promega) transfection reagent according to the manufacturer´s recommendations ratio (1 µg plasmid DNA, 3 µl FUGENE6) in OptiMEM (Gibco) and added to wells (Nunclon delta surface 6 well plates, Nunc) with 70−80% cell confluency in imaging buffer (DMEM without phenol red (Gibco), supplemented with glucose (4.5 g·L$^{-1}$), 10% v/v fetal bovine serum (ThermoFisher Scientific), 2 mM L-Glutamine (Gibco) and 1 mM sodium pyruvate (Gibco)) for 24−48 h. Transfected cells were then trypsinized, counted, and seeded into imaging plates (24 well glass bottom plate with high-performance #1.5 cover glass (Cellvis)). The labelling and imaging were conducted the following day.

#### Labelling and sample preparation

Cells were labelled with respective fluorophores (1 µM, 3 h at 37°C, in the dark) in imaging buffer and subsequently washed three to five times with prewarmed imaging buffer before fluorescence imaging.

#### Microscopy

Widefield imaging was performed at the Advanced Light Microscopy Facility (ALMF, EMBL Heidelberg), on a Nikon Ti-E microscope equipped with a Spectra X light engine (Lumencore) with a 40x objective (CFI P-Apo 40x Lambda/0.95/0.25−0.16) and imaged onto a scientific complementary metal-oxide-semiconductor camera (pco.edge 4.2 CL). The microscope was equipped with a $CO_2$ and temperature-controllable incubator (home-built, 37°C). A quad bandpass filter cube was used to image EGFP (excitation 485/20, emission 525/50), **JF$_{635}$-HTL/JF$_{630}$-HTL** (excitation 650/13, emission 692/40). Photoswitching was induced by illumination for 5 minutes (450 nm, 2.6 mW·cm$^{-2}$) using the custom-made LED box described above.

#### Image analysis

Statistical analysis as well as curve fitting was performed using GraphPad (Prism). All images were processed with ImageJ/Fiji and macros written therein unless otherwise stated.[18, 19] Segmentation of the nucleus was done in ImageJ/Fiji. Segmentation of actin was done using the Cellpose plugin for ImageJ/Fiji using the ‘cyto2’ model.[20] For the segmentation and evaluation of the psHaloTag1a$_{635}$ performance in mitochondria, Arif Khan of the bioimage analysis support team at EMBL wrote a specific Python script, which is based on cell segmentation of the cytosolic EGFP signal using a model trained on top of Cellpose model ‘cyto2’. The resulting cell labels were matched across images. The fluorescence intensity change $F/F_0$ was determined for each cell. Dead cells were excluded from the analysis. Arif Khan additionally provided a modified code to quantify intensity change in the photoswitching cycles of psHaloTag1a labelled with **JF$_{635}$-HTL** or **JF$_{630}$-HTL**, respectively.

To compare the basal brightness of the activated state, the fluorescence intensity of the psHaloTag1a$_{635/630}$ in the ON state was divided by the cytosolic GFP intensity. For actin, the same mask was used to obtain a value for the basal brightness of each cell. In contrast, for the nucleus and mitochondria, a mean EGFP value per field of view was calculated and the **JF$_{635}$** signal was divided by this value to account for differences in masks and areas, which led to larger variations.
